# Binding of synGAP to PDZ Domains of PSD-95 is Regulated by Phosphorylation and Shapes the Composition of the Postsynaptic Density

**DOI:** 10.1101/058016

**Authors:** Ward G. Walkup, Tara Mastro, Leslie T. Schenker, Jost Vielmetter, Rebecca Hu, Ariella Iancu, Meera Reghunathan, B. Dylan Bannon, B. Kennedy Mary

**Author notes:** To whom correspondence should be addressed: Mary B. Kennedy, Division of Biology and Biological Engineering 216-76, California Institute of Technology, Pasadena, CA, USA 91125.

## Abstract

SynGAP is a Ras/Rap GTPase-activating protein (GAP) present in high concentration in postsynaptic densities (PSDs) from mammalian forebrain where it binds to all three PDZ (<u>P</u>SD-95, <u>D</u>iscs-large, <u>Z</u>O-1) domains of PSD-95. We show that phosphorylation of synGAP by Ca^2+^/calmodulin-dependent protein kinase II (CaMKII) decreases its affinity for the PDZ domains as much as 10-fold, measured by surface plasmon resonance. SynGAP is abundant enough in postsynaptic densities (PSDs) to occupy about one third of the PDZ domains of PSD-95. Therefore, we hypothesize that phosphorylation by CaMKII reduces synGAP’s ability to restrict binding of other proteins to the PDZ domains of PSD-95. We support this hypothesis by showing that three critical postsynaptic signaling proteins that bind to the PDZ domains of PSD-95 are present at a higher ratio to PSD-95 in PSDs isolated from synGAP heterozygous mice.

## Introduction

The postsynaptic density (PSD) is an organized complex of signaling proteins attached to the postsynaptic membrane of excitatory glutamatergic synapses in the central nervous system. It comprises a network of scaffold proteins, the most prominent of which is PSD-95, a member of the MAGUK family (Membrane-Associated GUanylate Kinase-like proteins) (Kennedy, 2000). PSD-95 contains three PDZ domains that bind transmembrane receptors and cytosolic signaling proteins by attaching to four to seven residue sequences usually located at the C-terminus of the binding protein (Fig. 1A; Kornau et al., 1995; Kornau et al., 1997). PSD-95 associates with itself and other scaffold proteins via its guanylate kinase-like domain to form an interconnected network that spatially organizes receptors and biochemical signal transduction machinery at the postsynaptic site (Baron et al., 2006; Sheng and Kim, 2011).

**FIGURE 1.**
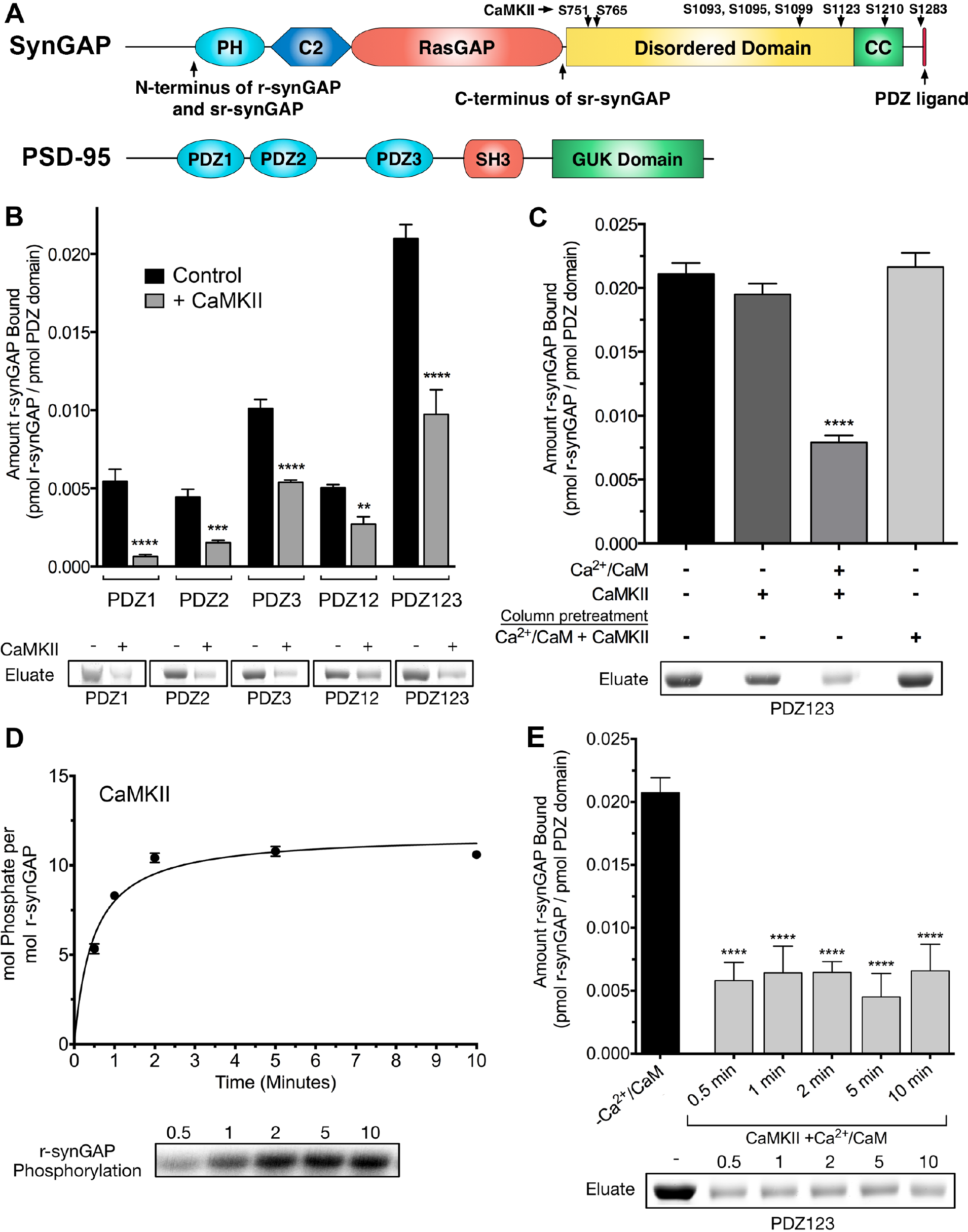
<u>Phosphorylation by CaMKII regulates association of r-synGAP with PDZ domains of PSD-</u> <u>95.</u> *A,* Domain diagrams of synGAP (Walkup IV et al., 2015) and PSD-95 (Cho et al., 1992). The boundaries of r-synGAP and sr-synGAP are indicated, as are the locations of the major sites phosphorylated by CaMKII, most of which are in the “disordered domain.” Numbering is based on rat isoform synGAP A1-α1. The 5 residue PDZ ligand is located at the C-terminus. The five major domains of PSD-95, including the approximate relationships of its three N-terminal PDZ domains are indicated. *B*, Association of r-synGAP with PDZ domains of PSD-95 before and after phosphorylation by CaMKII. R-synGAP was incubated in a phosphorylation mix for 10 min with either 0 CaMKII and 0 Ca^2+^/CaM (control) or 10 nM CaMKII and 0.7 mM CaCl_2_ /3.4 μM CaM (+ CaMKII) before binding to PDZ domain resins for 60 min at 25° C, as described under Materials and Methods. For comparison of binding of control to phosphorylated synGAP: PDZ1, p <0.0001; PDZ2, p = 0.0001; PDZ3, p <0.0001; PDZ12, p = 0.002; PDZ123, p <0.0001 *C,* Both Ca^2^ /CaM and CaMKII are required in the phosphorylation reaction to reduce binding of synGAP to PDZ123 resin. R-synGAP was incubated in the phosphorylation reaction without Ca^2+^/CaM or CaMKII (Control), with CaMKII alone, or with both before binding to PDZ resin. For comparison of Control to phosphorylation: + CaMKII, p = 0.2; +Ca^2+^/CaM, +CaMKII, p <0.0001. The final bar shows that phosphorylation of the PDZ123 domain resin itself doesn't alter binding of r-synGAP. PDZ123 domain affinity resin was phosphorylated for 60 min in the presence of CaMKII and 0.7 mM CaCl_2_/3.4 μM CaM before incubation with control r-synGAP (500 nM) for 60 min at 25° C. For comparison to Control, p = 0.7. *D,* Stoichiometry of phosphorylation of r-synGAP by CaMKII. R-synGAP (725 nM) was phosphorylated in the presence of CaMKII (10 nM), as described under “Materials and Methods.” At the indicated times, reactions were quenched by addition of 3x Laemmli sample buffer. Radiolabeled r-synGAP was isolated by SDS-PAGE and quantified as described under “Materials and Methods.” *E,* Change in affinity of r-synGAP for PDZ123 after phosphorylation by CaMKII for times corresponding to those measured in D. R-synGAP was phosphorylated for 0.5 to 10 min as described in *D* before incubation with PDZ123 domain affinity resin for 60 min as described under “Materials and Methods.” Control (-CaMKII,-Ca^2+^/CaM) is r-synGAP incubated in the phosphorylation reaction in the absence of CaMKII and Ca^2+^/CaM. For Comparison of Control to phosphorylation for 0.5, 1, 2, 5, or 10 min, p <0.0001. Pairwise comparisons among all phosphorylation conditions showed no significant differences (p from 0.4 to 1). Data shown in *B-E* are plotted as mean ± S.E. (n=4). For B, C and E, the statistical significance of differences in binding to PDZ domain resin relative to unphosphorylated r-synGAP control (-Ca^2+^/CaM) was determined by ordinary one way ANOVA (uncorrected Fisher’s LSD). ^**^, p<0.01; ^***^, p<0.001; ^#x002A;**^ *p*<0.0001.

Formation and modulation of excitatory synapses involves *de novo* assembly and/or re-arrangement of the proteins bound to PDZ domains of PSD-95 and other binding sites on the PSD scaffold (El-Husseini et al., 2000; Steiner et al., 2008; Sturgill et al., 2009). One class of glutamate receptor channels, NMDA-type glutamate receptors (NMDARs) bind directly to PDZ domains of PSD-95 via their cytosolic C-termini (Kornau et al., 1995). A second class, AMPARs, bind indirectly to PDZ domains of PSD-95 through two families of auxiliary subunits, TARPs (Transmembrane AMPA Receptor Regulatory Proteins) (Tomita et al., 2005) and LRRTMs (Leucine-Rich Repeat TransMembrane proteins) (de Wit et al., 2009). AMPARs, NMDARs, TARPs, and LRRTMs, along with the synaptic organizing molecule neuroligin, comprise the most highly enriched transmembrane proteins precipitated together with PSD-95 from the postsynaptic density fraction of mouse forebrain (Dosemeci et al., 2007). LRRTMs and neuroligin bind to presynaptic neurexin, which promotes formation of new presynaptic vesicle release sites (Siddiqui et al., 2010).

A smaller number of cytosolic signaling proteins also associate directly with PDZ domains of PSD-95 (Brenman et al., 1996; Chen et al., 1998; Kim et al., 1998; Murphy et al., 2006; Sakagami et al., 2008). Among these, the synaptic Ras/Rap GTPase activating protein synGAP is particularly interesting for several reasons. It is unusually abundant in the PSD fraction; indeed, it is nearly as abundant in PSD-95 complexes as PSD-95 itself (Chen et al., 1998; Cheng et al., 2006; Dosemeci et al., 2007). The *synGAP* gene displays a highly penetrant gene-dosage effect. Heterozygous deletion of synGAP in mice or humans produces a pronounced cognitive deficit (Komiyama et al., 2002; Hamdan et al., 2009; Hamdan et al., 2011; Berryer et al., 2013). This means that the amount of the synGAP protein is rate-limiting for at least one critical process at the synapse. Finally, the carboxyl tails of two prominent synGAP splice variants bind to all three of the PDZ domains of PSD-95 in yeast two hybrid assays (Kim et al., 1998). In contrast, most other PSD-95 binding proteins bind specifically either to PDZ domains 1 and 2, or to PDZ3 (e.g. Kornau et al., 1995; Irie et al., 1997). Taken together, these characteristics suggest that synGAP occupies a particularly large proportion of the PDZ domains of PSD-95 in most excitatory synapses; therefore, a 50% reduction in the amount of synGAP available to bind to PSD-95 might have pathological consequences for synaptic function.

Mutations in a single copy of synGAP have been causally implicated in ~ 9% of cases of non-syndromic cognitive impairment associated with either autism (ASD) or epilepsy (Berryer et al., 2013). Studies of the effects of such mutations in mice suggest a possible underlying neural mechanism.

Deletion of synGAP in cultured hippocampal neurons results in precocious formation of excitatory synapses. Enlarged postsynaptic spines containing clusters of PSD-95 develop in mutant neurons two days before they develop in wild-type neurons; and the precocious synapses contain a larger number of AMPARs (Vazquez et al., 2004). Early maturation of synapses in mice with a heterozygous deletion of synGAP shortens the critical period for experience-dependent synaptic development in the cortex (Clement et al., 2013). A corresponding shortening of the critical period in humans may underlie malformation of circuits with resulting cognitive impairment, autism, and/or epilepsy.

In mutant mouse neurons with a homozygous synGAP deletion, viral expression of recombinant wild-type synGAP reverses the precocious synapse formation (Vazquez et al., 2004). Spine heads enlarge and PSD-95 clusters enter synaptic spines at the normal time. In contrast, expression of synGAP lacking the five-residue C-terminal PDZ domain ligand (ΔSXV) does not reverse the phenotype. Indeed, we observed that, after expression of ΔSXV, the sizes of clusters of PSD-95 are larger than in the deletion mutant itself and continue to enter the spine earlier than in wild-type neurons. SynGAP bearing the ΔSXV mutation localizes normally to spines indicating that failure to rescue the phenotype is not caused by abnormal localization. This observation led us to hypothesize that binding of synGAP to the PDZ domains of PSD-95 might normally restrict the binding of other proteins that influence clustering of PSD-95 and its movement into the spine. Recent work suggests that those proteins could include neuroligins (Sudhof, 2008), LRRTMs (Siddiqui et al., 2010) or TARPs (Tomita et al., 2005).

Phosphorylation of synGAP by CaMKII has recently been shown to induce movement of synGAP out of the PSD in living neurons (Yang et al., 2013; Araki et al., 2015). In our recent study of phosphorylation sites for CaMKII on r-synGAP, a soluble recombinant form of synGAP missing the first 100 N-terminal residues, we found one site (S1283) located very near the C-terminal PDZ domain ligand (Fig. 1A; Walkup IV et al., 2015). Therefore, we tested whether phosphorylation of this site or others by CaMKII reduces the affinity of synGAP for PDZ domains of PSD-95. Here we demonstrate that phosphorylation of several sites in the disordered domain of synGAP and of S1283 dramatically reduces binding of synGAP to all of the PDZ domains on PSD-95. We also report the unexpected finding that the disordered domain of synGAP binds Ca^2+^/CaM with high affinity and this binding causes a smaller reduction of affinity of synGAP for PDZ3. We hypothesize that phosphorylation of synGAP in the PSD, triggered by activation of NMDARs, could substantially reduce the ability of synGAP to compete with other proteins for binding to the PDZ domains of PSD-95, permitting increased equilibrium binding of neuroligin, TARPs and/or LRRTMs. We support the hypothesis that binding of synGAP to PSD-95 restricts binding of other proteins by showing that the number of TARPs, LRRTM2, and neuroligin-2 molecules per molecule of PSD-95 is significantly increased in PSD fractions isolated from mice with a heterozygous deletion of synGAP compared to those isolated from wild type mice.

We propose that reconfiguration of the PSD triggered by phosphorylation of synGAP and the resulting decrease in its affinity for PDZ domains of PSD-95 facilitates formation of new synapses and/or leads to strengthening of existing synapses in part by allowing increased trapping of AMPARs in the PSD during the early stages of induction of long-term potentiation (LTP; Opazo and Choquet, 2011). This proposed mechanism is consistent with the “slot” hypothesis for addition of new AMPA-receptors during induction of LTP (Shi et al., 2001; Henley and Wilkinson, 2016), and we discuss how it can account for many previous experimental observations.

## Results

*Phosphorylation of R-synGAP by CaMKII Reduces Its Binding to PDZ Domains of PSD-95–* SynGAP can be expressed in bacteria and purified in a soluble form by deleting the first 102 residues of its N-terminus (Walkup IV et al., 2015). This version of synGAP, termed r-synGAP, retains all of the identified functional domains, the “disordered” regulatory domain, and the C-terminal PDZ ligand (Fig. 1A). In a previous study we showed that r-synGAP is phosphorylated by CaMKII at several residues including S1283, which is 7 residues upstream of the PDZ domain ligand located at residues 1290-1293 (Walkup IV et al., 2015). Because this phosphorylation site is so near the PDZ ligand, we wondered whether its phosphorylation, or phosphorylation of other sites by CaMKII, would interfere with binding of synGAP to PDZ domains of PSD-95. To test this, we incubated r-synGAP with affinity resins substituted with recombinant PDZ domains as described under Materials and Methods. The beads contained PDZ1 (61-151), PDZ2 (155-249), PDZ3 (302-402), a fragment containing PDZ1 and PDZ2 (PDZ12, 61-249), or a fragment containing all three PDZ domains (PDZ123, 61-402). Binding of r-synGAP to the beads was tested with or without a prior 10 min phosphorylation by CaMKII. As expected, without phosphorylation, r-synGAP binds specifically to each of the three PDZ domains (Fig. 1B). In this assay, its binding is highest to PDZ3. Binding of r-synGAP to PDZ123 reveals a substantial avidity effect; that is, the amount bound per individual PDZ domain is twice that bound to PDZ3 alone and four times that bound to either PDZ1 or PDZ2 alone.

Phosphorylation by CaMKII reduces binding of synGAP to all of the individual PDZ domains and to to the constructions comprising PDZ12 and PDZ123 (Fig. 1B). The reduction in binding requires the presence of both Ca^2+^/CaM and CaMKII in the phosphorylation reaction mixture (Fig. 1C). The fourth bar of Fig. 1C shows that the reduction in binding is not caused by phosphorylation of PDZ domains on the column by residual CaMKII. We have shown previously that as many as 10 sites on synGAP are phosphorylated by CaMKII (Walkup IV et al., 2015). Approximately 5 of these, including site S1123, are fully phosphorylated after a 0.5 min reaction (Walkup IV et al., 2015 and Fig. 1D). To test whether the reduction in binding depends primarily on phosphorylation of the rapidly phosphorylated sites or requires phosphorylation of most of the sites, we tested binding of r-synGAP to PDZ123 after phosphorylation for various times (Fig. 1E). The reduction in binding is maximal after 0.5 min, indicating that phosphorylation of the more rapidly phosphorylated sites is sufficient for full reduction of affinity.

*Affinity of R-synGAP for the PDZ Domains of PSD-95 Determined by Surface Plasmon Resonance (SPR)–* The three PDZ domains of PSD-95 are located in the N-terminal half of the protein from residues 61 to 402. The first two PDZ domains are separated by 4 residues and the third is 53 residues downstream of PDZ2. We determined the affinities of r-synGAP for individual PDZ domains, for PDZ12 and for PDZ123 by a “competition in solution” assay in which SPR is used to detect the amount of free r-synGAP in solutions containing a constant amount of r-synGAP and varying amounts of recombinant PDZ domains (Nieba et al., 1996; Lazar et al., 2006; Abdiche et al., 2008). To detect the free synGAP, recombinant PDZ domains are immobilized on the Biacore chip as described under Materials and Methods. We used the competition method rather than conventional Biacore measurements in which varying concentrations of r-synGAP are applied to a chip containing immobilized PDZ domains because concentrations of r-synGAP above ~100 nM produced a large bulk resonance signal caused by high viscosity that obscured the change in resonance produced by its binding to PDZ domains. The competition assay eliminates the need to apply high concentrations of synGAP to the chip.

We generated a standard curve in which the maximum resonance responses of a series of concentrations of synGAP from 0 nM to 50 nM (Fig. 2A, grey traces) were determined and plotted against synGAP concentration (Fig. 2B, large grey dots). The data were fit with a hyperbolic curve. The maximum resonance response of a series of mixtures containing 25 nM r-synGAP and increasing concentrations of PDZ1 from 0 nM to 10 μM were measured, and the concentration of r-synGAP remaining free to bind to PDZ1 on the chip was then determined from the standard curve (Fig. 2B, small black dots). A K_D_ of 140 ± 30 nM (Table 1) was calculated as described under Materials and Methods (Fig. 2C). We used the same method to measure K_D_s for PDZ2 and PDZ3 (Fig. 3A and B, respectively) and K_Dapp_s for PDZ12 and PDZ123 (Fig. 3C and D, respectively). The values are summarized in Table 1. We obtained an additional value of 730 ± 50 nM for the K_D_ of PDZ3 by a conventional Biacore assay, which is in good agreement with the K_D_ measured by the competition assay. These data show that, under these conditions, PDZ1 has a higher affinity for synGAP than does PDZ3.

**Table 1.** <u>Affinities of R-synGAP for PDZ Domains of PSD-95</u> Dissociation constants (K_D_ or K_Dapp_) for the interactions of synGAP with PDZ domains of PSD-95 were determined by the Biacore/SPR “competition in solution” method as described under “Materials and Methods.” In one experiment, the K_D_ for PDZ3 was determined by conventional SPR as described under “Materials and Methods.” Goodness of Fit refers to the fit of the data shown in Figs. 2, 3, and 5 to the equation relating synGAP_free_ to PDZ domain concentration described under “Materials and Methods.” Data are expressed as mean ± S.E.

| PDZ Domain from PSD-95 | No. of Experiments | Dissociation Constant ($K_D$) for Binding r-synGAP (nM) | Goodness of Fit ($R^2$) |
| --- | --- | --- | --- |
| PDZ1 | 3 | $220 \pm 30$ | 0.908 - 0.947 |
| PDZ2 | 2 | $1500 \pm 100$ | 0.967, 0.969 |
| PDZ3 | 2 | $620 \pm 70$ | 0.951, 0.9624 |
| PDZ3 | 1 | $730 \pm 50$ (by conventional SPR) | N.A. |
|  |  | **Apparent Dissociation Constant ($K_{Dapp}$) for Binding to R-synGAP (nM)** |  |
| PDZ12 | 4 | $350 \pm 40$ | 0.931 - 0.987 |
| PDZ123 | 6 | $4.7 \pm 0.6$ | 0.957 - 0.985 |
| PDZ123 | 3 | (CaMKII Phosphorylated r-synGAP)<br>$46 \pm 10$ | 0.810 - 0.880 |
| PDZ123 | 2 | (S1283D r-synGAP)<br>$16 \pm 3$ | 0.953, 0.954 |

**FIGURE 2.**
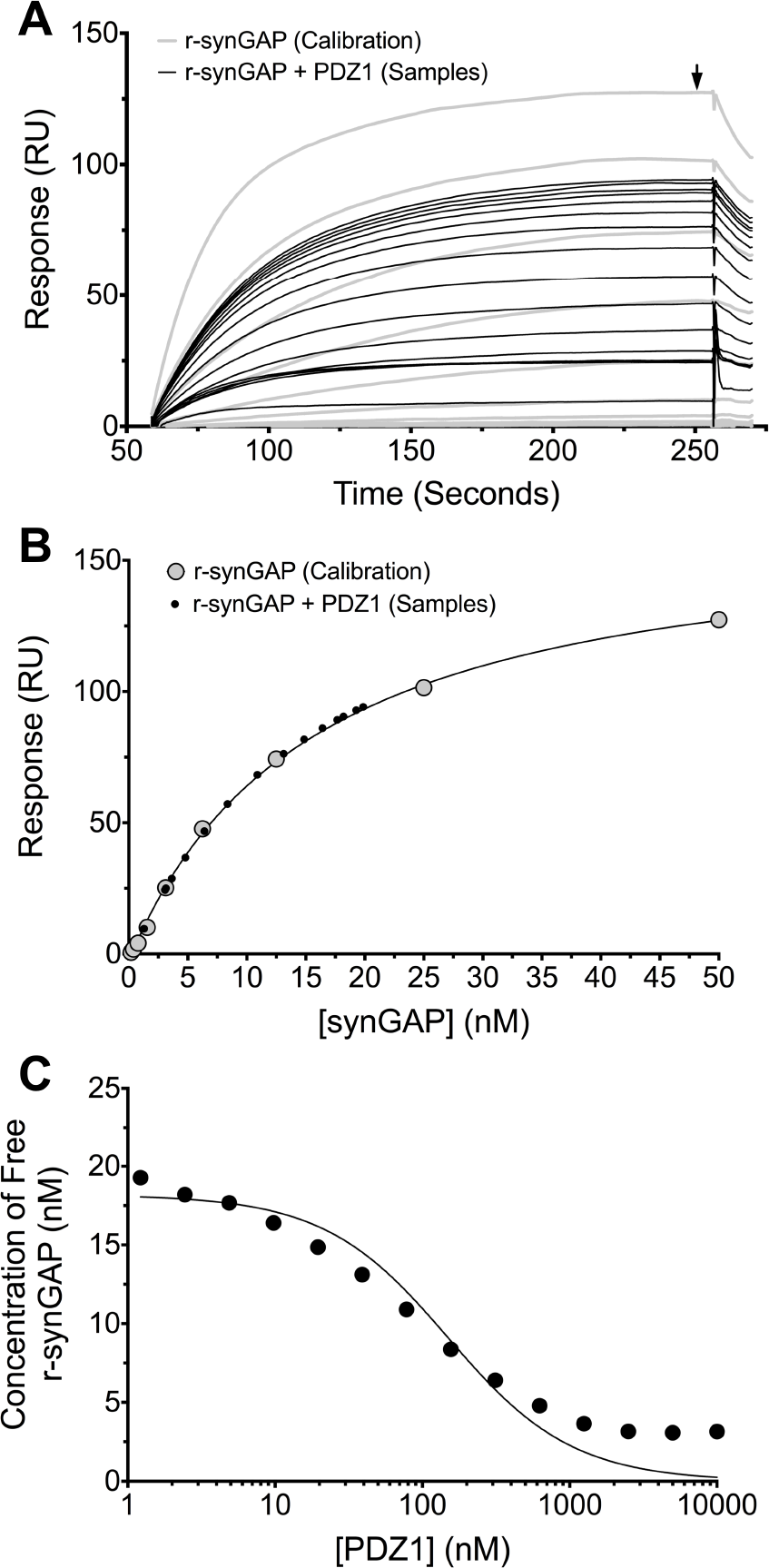
<u>Measurement of affinity of r-synGAP for PDZ1 of PSD-95 by the “competition in solution”</u> <u>method.</u> *A,* Biacore sensorgrams showing the calibration curves (grey lines) for binding of 0-50 nM r-synGAP and the measurement of free r-synGAP (samples; black lines) in mixtures containing 25 nM r-synGAP and 0-10 <M PDZ1 domain. Free r-synGAP was detected by binding to PDZ1 domains immobilized on a Biacore chip as described under Materials and Methods. *B,* A standard calibration curve was constructed by plotting the maximum calibrated resonance responses (marked by arrow in A) against the corresponding concentrations of r-synGAP (large grey dots). The maximum resonance responses for each sample mixture were plotted on the standard curve to determine the free r-synGAP concentrations in each mixture (black dots). *C,* Plot of free r-synGAP concentrations determined in *B* against the log of PDZ domain concentrations (black circles). The data were fit to the binding equation shown in Materials and Methods with the use of Graphpad Prism software. A K_D_ value (Table 1) was calculated from the equation as described under Materials and Methods.

**FIGURE 3.**
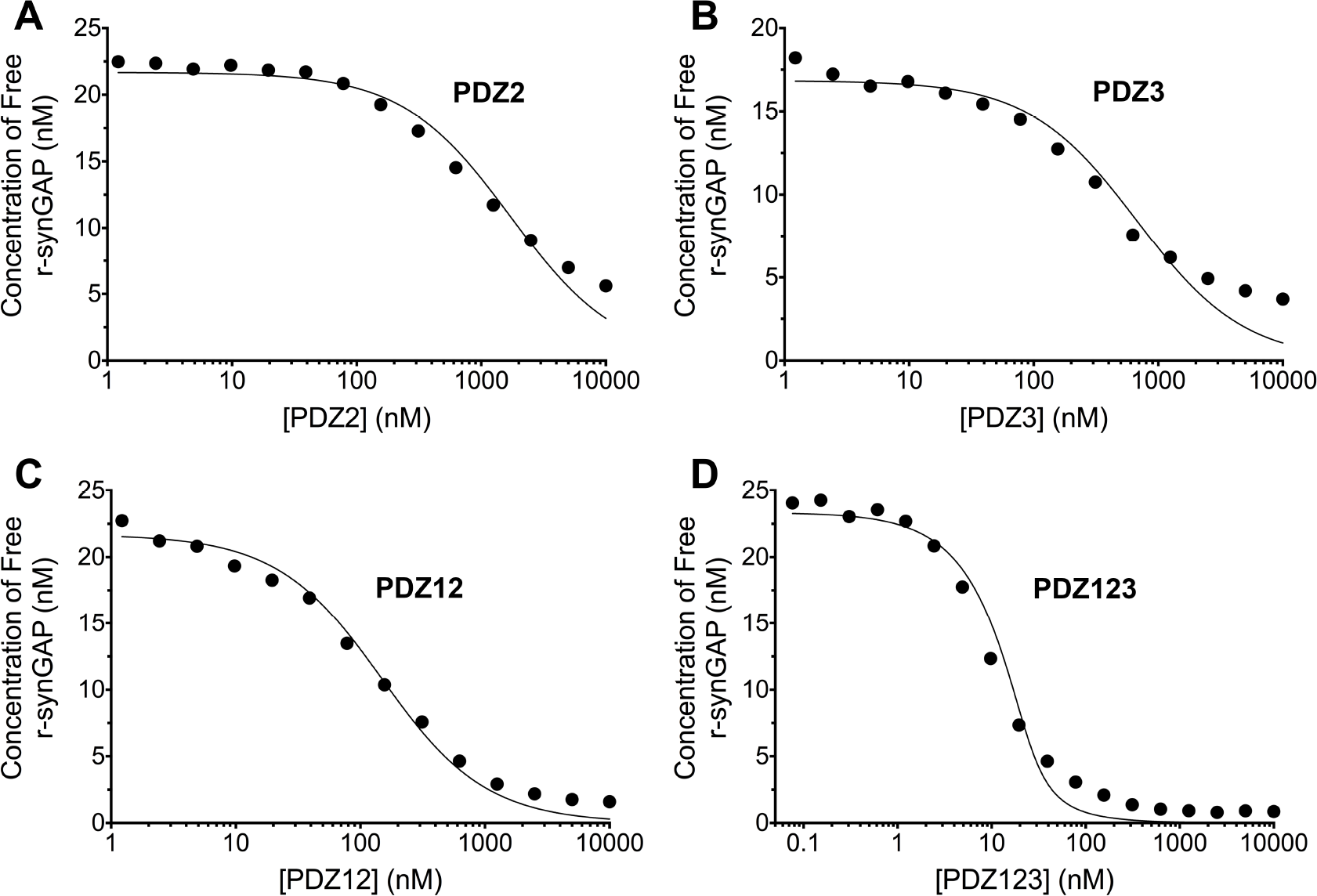
<u>Affinities and apparent affinities of r-synGAP for PDZ2, PDZ3, PDZ12 and PDZ123</u> <u>domains of PSD-95 determined by the “competition in solution” method.</u> The concentrations of free r-synGAP in sample mixtures containing each of the indicated PDZ domains were measured as described in Fig. 3A and B, and under Materials and Methods. The values (black dots) were plotted against the log of the PDZ domain concentration and fit to a binding curve as described in Fig. 3C. Representative experiment for *A,* PDZ2; *B,* PDZ3; *C,* PDZ12; and *D,* PDZ123. The calculated K_D_ and K_Dapp_ values from all experiments are listed in Table 1.

*Binding of R-SynGAP Phosphosite Mutants to PDZ123 Affinity Resin–* To determine which phosphorylation sites are important for the reduction in binding to PDZ domains that we observed in Fig. 1A, we first examined binding of recombinant mutants of synGAP to PDZ123 affinity resin. Neither mutation of site S1123 to alanine or aspartate nor double mutation of sites S1093 and S1123 to alanine alters binding of r-synGAP to PDZ123 before phosphorylation (Fig. 4A). However, each of these mutations reduces the effect of phosphorylation on binding compared to wild-type after 0.5 min of phosphorylation by CaMKII. Mutation of S1123 to alanine had the same effect as double mutation of S1093 and S1123 to alanines (p = 0.6), indicating that phosphorylation of S1093 has relatively little effect on binding to PDZ domains; whereas, phosphorylation of S1123 contributes to the reduction of binding to PDZ domains, but is not sufficient to produce the maximum reduction of binding. In contrast to S1123, mutation of S1283 alone to aspartate reduces the binding of r-synGAP to PDZ123 by ~50% relative to wild-type before phosphorylation (Fig. 4B), suggesting that phosphorylation of this site alone causes substantial loss of affinity for PDZ domains. Notably, none of the mutations interfere with the effect of ten min of phosphorylation (Fig. 4A and B). Taken together, these results mean that phosphorylation of S1283 alone significantly reduces binding of synGAP to PDZ domains, however maximum loss of binding can be accomplished only by cumulative phosphorylation over ten min of several sites within the regulatory disordered domain (See Fig. 1C in Ref. Walkup IV et al., 2015).

**FIGURE 4.**
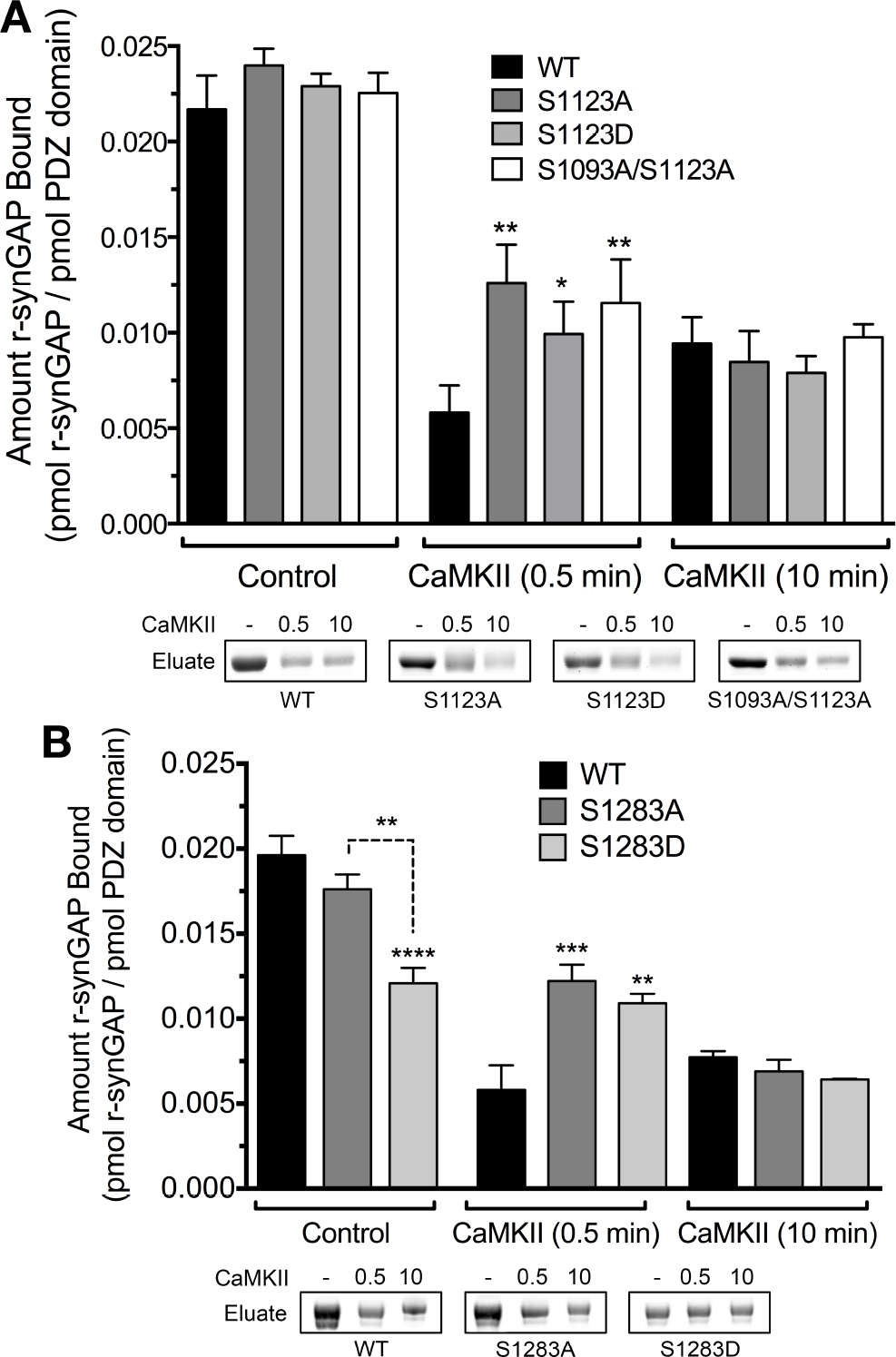
<u>Effect of phosphorylation by CaMKII on association of PDZ123 domains with phospho-</u> <u>deficient and phospho-mimetic mutants of r-synGAP.</u> Wild-type and mutant r-synGAP were incubated with phosphorylation mixtures for 10 min without (Control) or for 0.5 or 10 min with 10 nM CaMKII, 0.7 mM CaCl_2_/3.4 <M CaM (CaMKII) then incubated with PDZ123 affinity resin for 60 min at 25° C, as described under “Materials and Methods.” *A,* Binding of r-synGAP and r-synGAP mutants S1123A, S1123D, and S1099A/S1123A are represented by the indicated shades of grey bars. For comparison of WT to mutants after 0.5 min phosphorylation: S1123A, p = 0.001; S1123D, p = 0.04; S1093A/S1123A, p = 0.005. For comparison of WT to mutants after 10 min phosphorylation: S1123A, p = 0.625; S1123D, p = 0.437; S1093A/S1123A, p = 0.867. For comparison of WT and each mutant after 0.5 and 10 min phosphorylation: WT, p = 0.07; S1123A, p = 0.04; S1123D, p = 0.30; S1093A/S1123A, p = 0.366. *B,* Binding of r-synGAP and r-synGAP mutants S1283A and S1283D are represented by the indicated shades of grey bars. For comparison of WT to mutants before phosphorylation: S1283A, p = 0.2; S1283D, p <0.0001; For comparison of S1283A to S1283D, p = 0.0009. For comparison of WT to mutants after 0.5 min phosphorylation: S1283A, p = 0.0001; S1283D, p = 0.002. For comparison of WT and mutants after 0.5 and 10 min phosphorylation: WT, p = 0.2, S1283A, p = 0.001; S1283D, p = 0.007. Data are mean ± S.E. (n=4). The statistical significance of differences in PDZ domain binding among wild-type and mutant samples was determined by ordinary one way ANOVA (uncorrected Fisher’s LSD). *\*, p*<0.05; ^**^, *p*<0.01; ^***^, p<0.005;^****^, *p*<0.0001.

*Effect of Phosphorylation on Affinity of R-synGAP for PDZ123 Measured by SPR–* We measured the apparent dissociation constant (K_Dapp_) for binding to PDZ123 of r-synGAP phosphorylated for 10 min by CaMKII, and of the phosphomimetic mutant r-synGAP S1283D. Phosphorylation for 10 min by CaMKII increases the K_Dapp_ of r-synGAP approximately ten-fold (Fig. 5A, Table 1); whereas mutation of S1283 to aspartate increases the K_Dapp_ approximately four-fold (Fig. 5B, Table 1). Thus, cumulative phosphorylation of several sites on r-synGAP can reduce affinity for PDZ domains by an order of magnitude; whereas, addition of a negative charge at S1283 alone can reduce the affinity by a factor of four.

**FIGURE 5.**
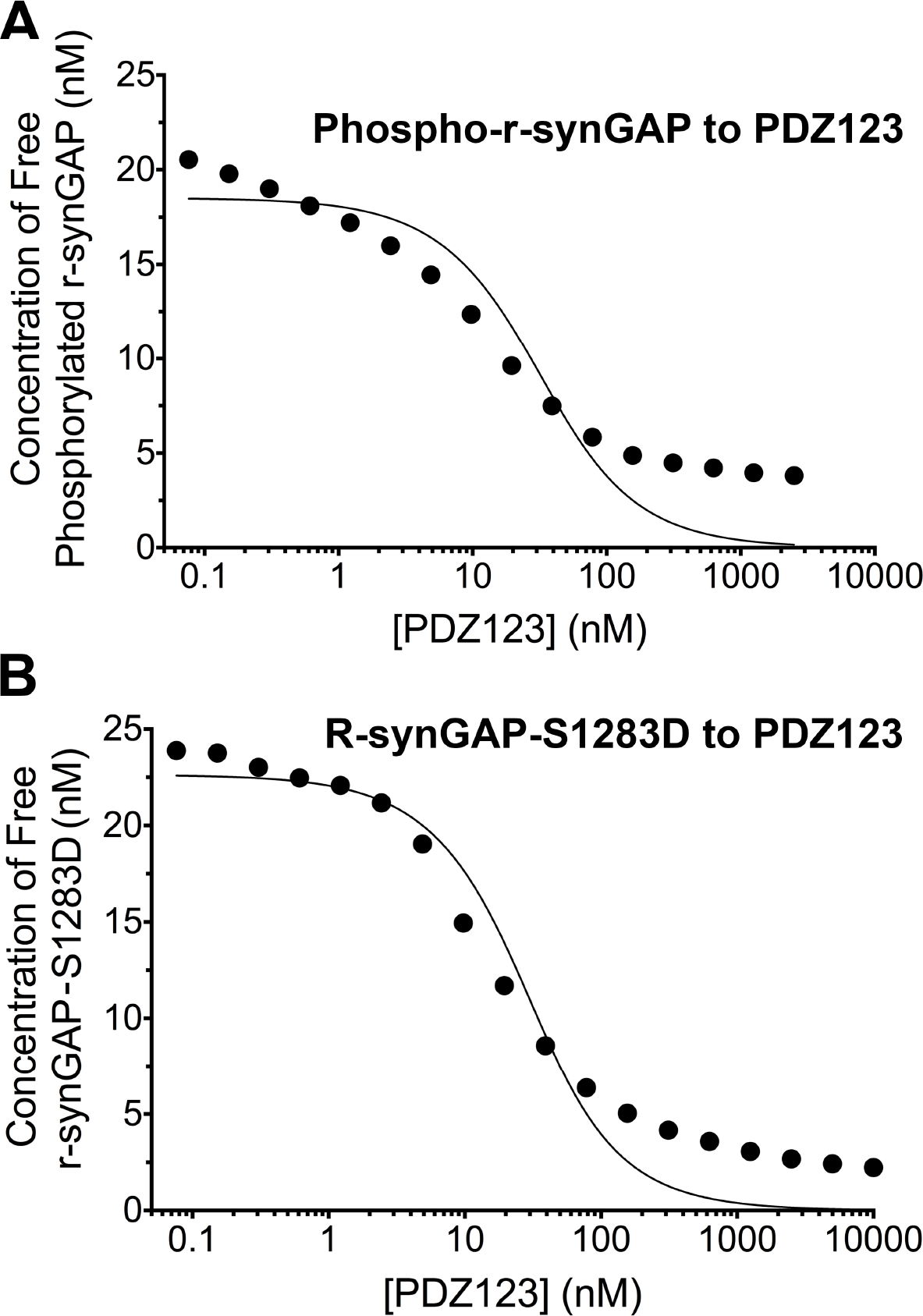
<u>Apparent affinities of phosphorylated r-synGAP and r-synGAP-S1283D for PDZ123</u> <u>determined by the “competition in solution” method.</u> Representative plots of the concentrations of (A) free phospho-r-synGAP phosphorylated as described for PDZ Binding Assays under Materials and Methods, and (B) r-synGAP-S1283D, measured in sample mixtures containing PDZ123 as described in Fig. 3A and B, and under Materials and Methods. The values (black dots) were plotted against the log of the PDZ123 concentration in the mixture and fit to a binding curve as described in Fig. 3C. The calculated K_Dapp_ values are listed in Table 1.

*Ca^2+^/CaMBinds Directly to R-synGAP–* While studying phosphorylation of r-synGAP by CDK5 (Walkup IV et al., 2015), we found that the presence of Ca^2+^/CaM in reactions with either CDK5/p35 or CDK5/p25 doubled the rate and stoichiometry of the phosphorylation (Fig. 6A and B). Inclusion of Ca^2+^ or CaM alone in the phosphorylation reactions did not alter the rates or stoichiometry.

**FIGURE 6.**
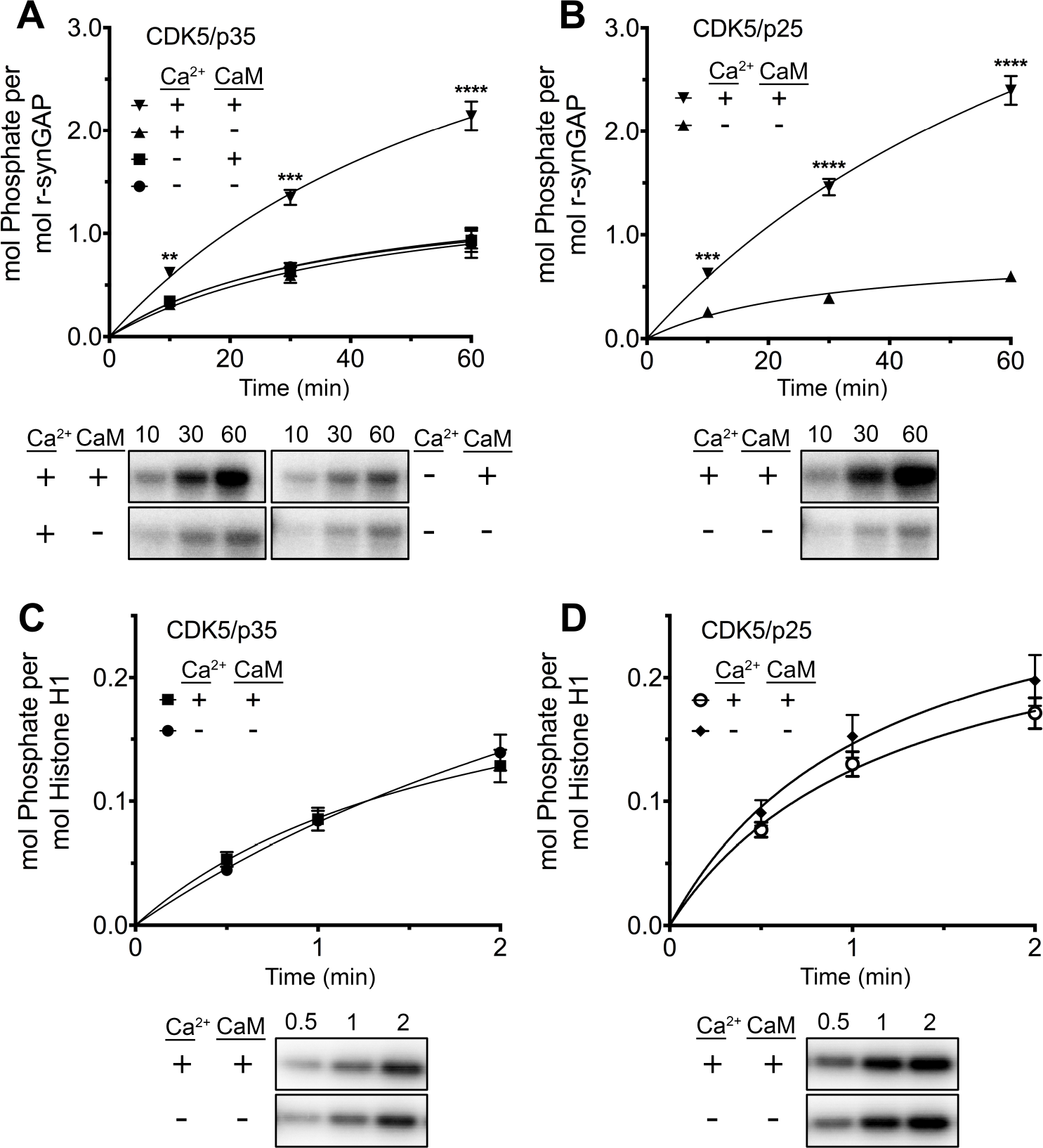
<u>Effect of Ca^2+^/CaM on stoichiometry of phosphorylation of r-synGAP and Histone H1 by CDK5.</u> Stoichiometry of phosphorylation of r-synGAP *(A-B)* and Histone H1 *(C-D)* by CDK5/p35 or CDK5/p25. R-synGAP (286 nM) or Histone H1 (4.3 μM) were incubated with CDK5/p35 or CDK5/p25 as described uner “Materials and Methods” in the presence or absence of 0.7 mM CaCl_2_ or 3.4 CaM, as indicated in each panel. Reactions were quenched at the indicated times by addition of 3x Laemmli sample buffer and radiolabeled r-synGAP and Histone H1 were quantified as described under Materials and Methods. For comparison of phosphorylation in the presence and absence of Ca^2+^/CaM: *A)* 10 min, p = 0.01; 30 min, p <0.0001; 60 min, p <0.0001. *B)* 10 min, p = 0.001; 30 min, p <0.0001, 60 min, p <0.0001. *C)* 0.5 min, p = 0.5; 1 min, p = 0.9; 2 min, p = 0.4. *D)* 0.5 min, p = 0.5; 1 min, p = 0.3; 2 min, p = 0.2. Data are plotted as mean ± S.E. (n = 4-7). The statistical significance of differences in phosphorylation in the presence of Ca^2+^ and CaM were determined by ordinary one way ANOVA (uncorrected Fisher's LSD). *\*\*, p*<0.01; ^***^, *p*<0.001; ^****^, *p*<0.0001.

**FIGURE 6-figure supplement 1.**
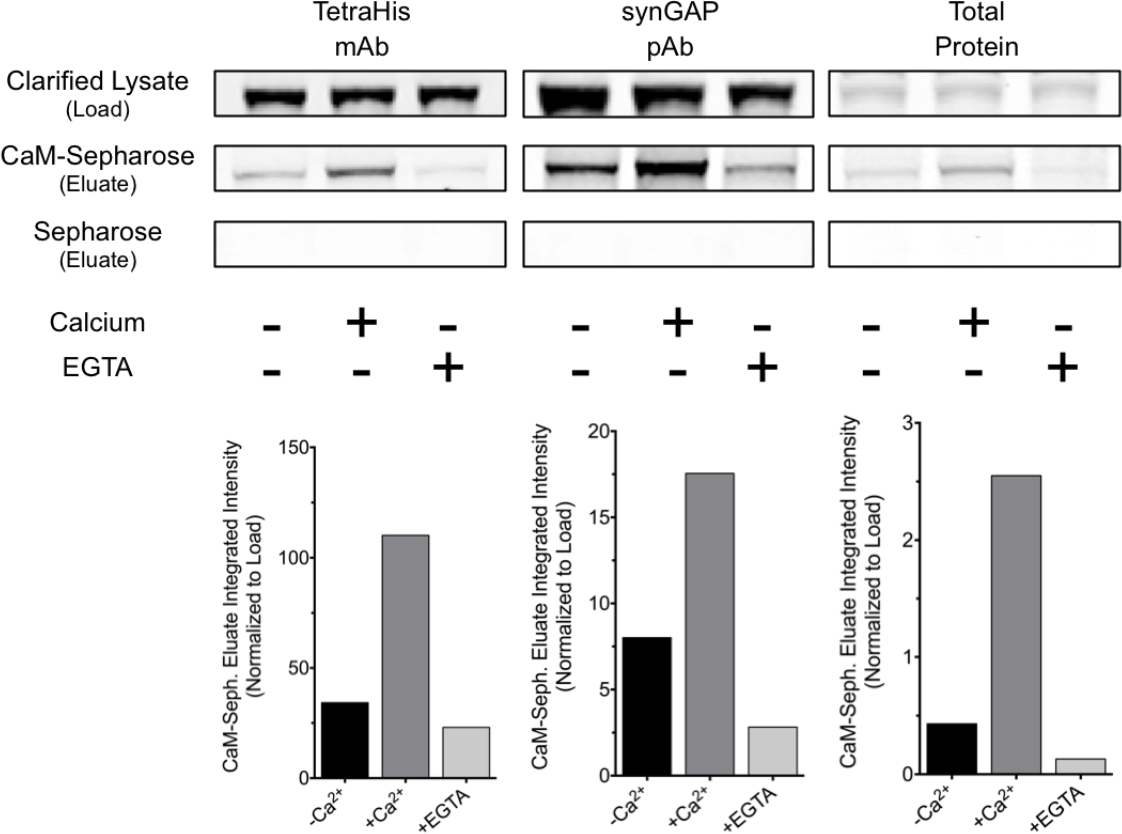
<u>R-synGAP binds to CaM affinity resin.</u> Clarified *E. coli* lysate (Load) containing r-synGAP was incubated with CaM-Sepharose 4B or control Sepharose 4B resin in the presence of 0 or 5 mM CaCl2 and 0 or 10 mM EGTA, as described under “Materials and Methods.” After washing, bound protein was eluted from the resin with 100 mM EGTA (Eluate), fractionated by SDS-PAGE, and visualized by staining with Gel Code Blue (Total Protein) or transferred to a PVDF membrane. R-synGAP was detected on the immunoblots with anti-synGAP or anti-TetraHis antibodies, as described under “Materials and Methods.” In the absence of exogenous calcium, r-synGAP bound weakly to the CaM-Sepharose, but not to control Sepharose beads. When 5 mM Ca^2+^ was included in the binding and wash buffers its binding to CaM-Sepharose increased, while addition of 10 mM EGTA to the buffers nearly abolished binding.

**Figure 6-figure supplement 2.**
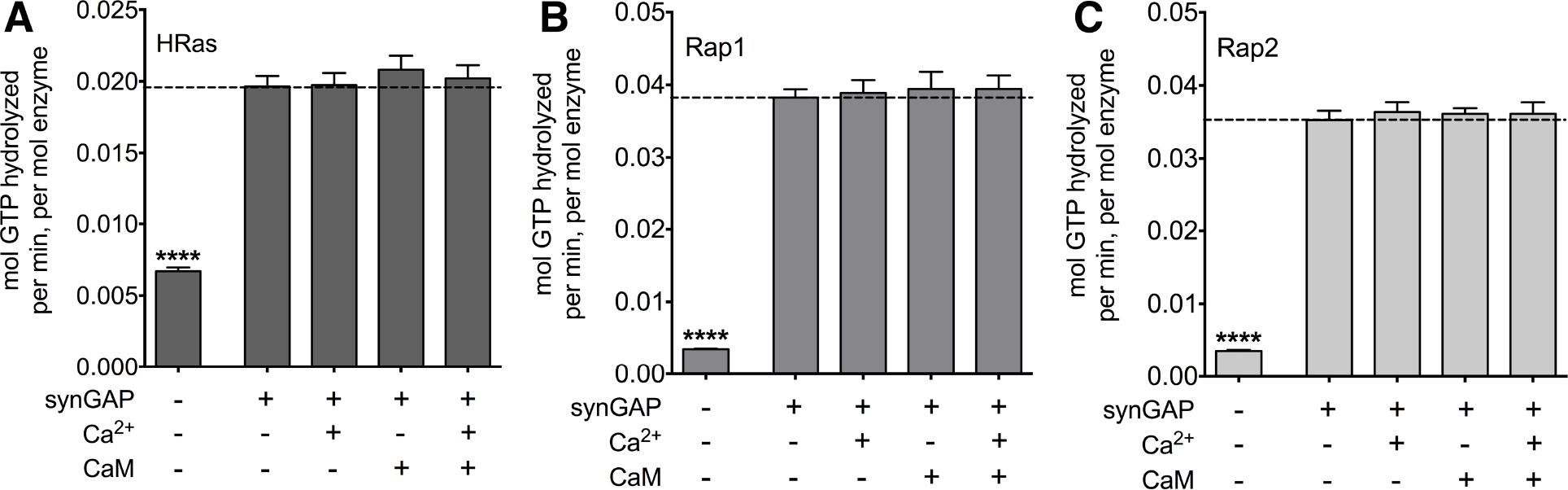
<u>Effect of binding of Ca^2+^/CaM on GAP activity of r-synGAP.</u> The GAP activity of r-synGAP (250 nM) for the indicated GTP-binding protein was assayed as described in Walkup et al. (2015) except that 1 mM Ca^2+^, 3.4 μM CaM, or both were added to the GAP assay as indicated. *A*, HRas GAP activity; *B*, Rap1 GAP activity; and *C*. Rap2 GAP activity. Data are mean ± S.E. The statistical significance of differences between activity in the presence of r-synGAP in the absence of Ca^2+^ or CaM (second column) and all other conditions was determined by ordinary one way ANOVA (uncorrected Fisher’s LSD). ^****^, p<0.0001. For comparison of GAP activities in the absence and presence of synGAP: HRas, Rap1 or Rap2, p < 0.0001. For comparisons of reactions containing synGAP: HRas, p ranged from 0.3 to 0.9; Rap1, p ranged from 0.5 to 0.7; Rap2, p ranged from 0.5 to 0.6.

We tested whether this effect resulted from binding of Ca^2+^/CaM to CDK5/p35 (e.g. He et al., 2008) by comparing the rates of phosphorylation of histone H1, a well-known substrate of CDK5 in the presence and absence of Ca^2+^/CaM (Fig. 6C and D). Phosphorylation of Histone H1 by either CDK5/p35 or CDK5/p25 was unaffected by Ca^2+^/CaM. This result suggests that Ca^2+^/CaM binds directly to r-synGAP, causing a substrate-directed enhancement of its phosphorylation.

To further verify that Ca^2+^/CaM binds directly to r-synGAP, we showed that r-synGAP binds to a CaM-Sepharose affinity resin in a Ca^2+^-dependent manner, as would be expected if it binds Ca^2+^/CaM specifically and with significant affinity (Fig. 6–figure supplement 1). We found that the presence of Ca^2+^/CaM alone in a Ras-or Rap-GAP assay has no effect on the GAP activity of synGAP (Fig. 6–figure supplement 2).

*Affinity of Binding of R-synGAP to Ca^2+^/CaM–* We measured the affinity of binding of Ca^2^ /CaM to r-synGAP by the conventional SPR method on the Biacore as described under Materials and Methods. CaM was immobilized on a chip, and r-synGAP was applied to it at concentrations from 0 to 75 nM (Fig. 7A). Analysis of the equilibrium phase of association at each concentration (Fig. 7B) yielded a K_D_ of 9 ± 1 nM, indicative of high affinity binding.

**FIGURE 7.**
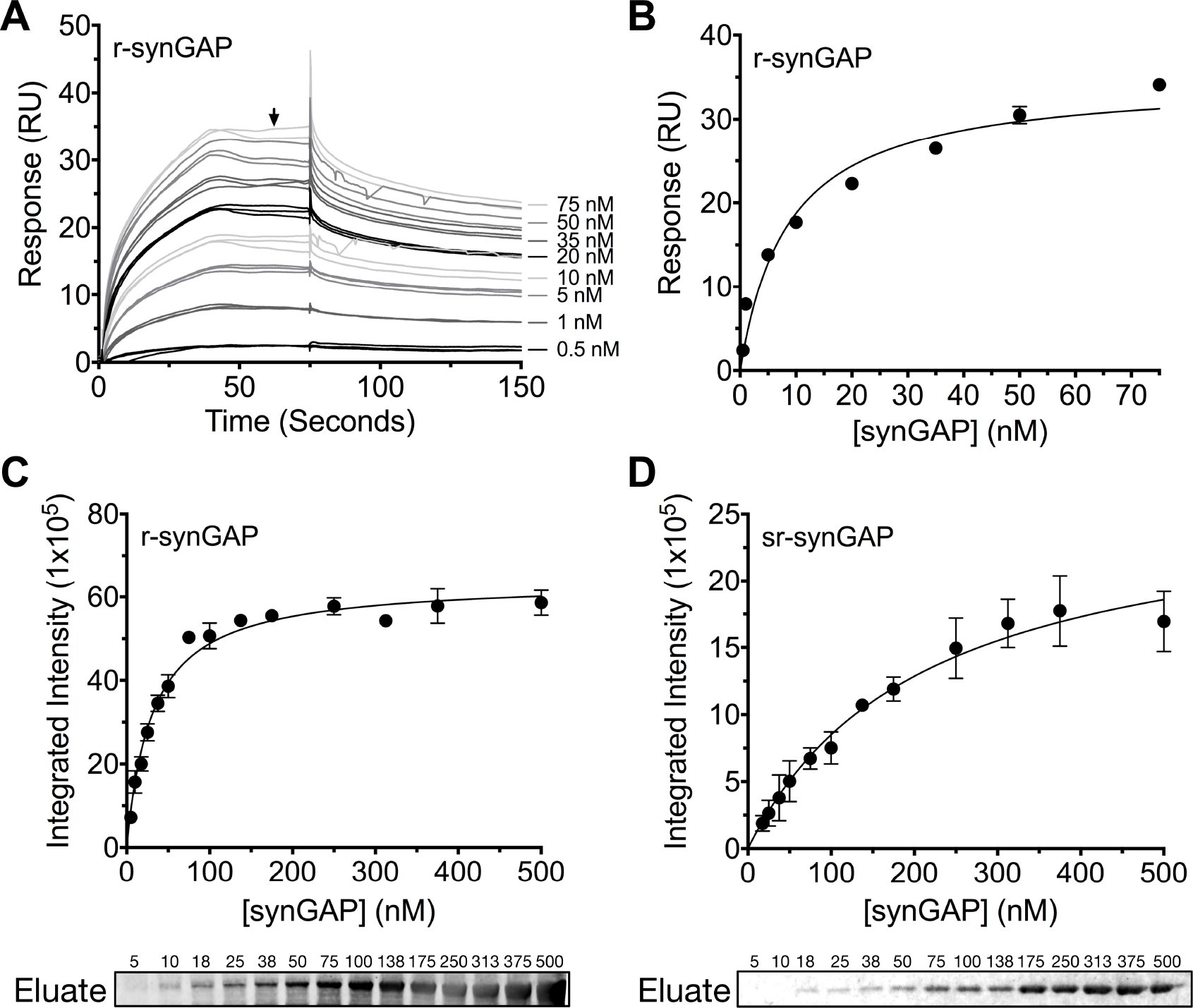
<u>Affinity of r-synGAP for Ca^2+^/CaM determined by equilibrium analysis.</u> (*A-B*) The affinity of r-synGAP for Ca^2+^/CaM was measured by SPR with CaM immobilized on the chip and r-synGAP injected at 0-50 nM onto the chip surface as described under Materials and Methods. *A,* Sensorgrams with the blank and reference flow cell readings subtracted show the response upon injection of r-synGAP onto the chip surface (0-75 seconds) and its dissociation from the chip surface (75-150 seconds). *B,* RUs at equilibrium (marked by arrow in A) were plotted against the corresponding concentrations of r-synGAP and fitted to a hyperbolic curve. A K_D_ of 9 ± 1 nM was calculated as described under Materials and Methods. *C and* D, The affinities of r-synGAP and sr-synGAP (0-500 nM) for Ca^2+^/CaM were measured by incubation with CaM-Sepharose resin as described under Materials and Methods. Integrated intensities of bound r-and sr-synGAP were measured from immunoblots as described under Materials and Methods and plotted versus the corresponding concentrations incubated with resin. Integrated intensities from Western blots were linear over the range of r-and sr-synGAP concentrations used in the assays *C,* r-synGAP; and *D,* sr-synGAP. Data in *C* and *D* are plotted as mean ± S.E. (n = 3).

To begin to define the location of the high affinity Ca^2+^/CaM binding site, we compared the affinities for Ca^2+^/CaM of r-synGAP and a C-terminal truncated protein, sr-synGAP (residues 103-725; Fig. 1A) by a bead-binding assay as described under Materials and Methods. We measured the amount of each protein bound to a fixed amount of CaM-Sepharose after incubation with increasing concentrations (Fig. 7C and D). Both r-synGAP (Fig. 7C) and sr-synGAP (Fig. 7D) showed saturable binding to the CaM-Sepharose resin, and did not bind to control Sepharose beads lacking CaM (data not shown). The data were fit to hyperbolic curves and the K_D_’s for binding of r-synGAP and sr-synGAP to Ca^2+^/CaM were calculated to be 31 ± 3 nM and 210 ± 30 nM, respectively. Thus, the high affinity site appears to be located in the regulatory disordered region of r-synGAP, which is missing in sr-synGAP. The K_D_’s determined by the bead-binding assay (31 ± 3 nM) and Biacore equilibrium binding (9 ± 1 nM) are in the range of those reported for calcineurin (PP2B) and CaMKII, 1-10 nM (Hubbard and Klee, 1987; Cohen and Klee, 1988) and 40-80 nM (Miller and Kennedy, 1985; Meyer et al., 1992; Hudmon and Schulman, 2002), respectively. We did not detect any binding when sr-synGAP was injected onto the CaM-substituted Biacore chip at concentrations from 10-2500 nM. Thus, the relatively weak binding of sr-synGAP observed in the bead-binding assay is not reproducible when measured on the Biacore chip. These data indicate that Ca^2+^/CaM binds only weakly, if at all, to the N-terminal half of synGAP. A meta-analysis algorithm for detecting potential CaM-binding domains (Mruk et al., 2014) predicts two Ca^2+^/CaM binding sites in the C-terminal half of synGAP, one from residues 1000-1030 and another in the putative coiled coil domain from residues 1229-1253. The SPR measurements do not allow us to confirm or to rule out the presence of two high affinity sites of similar affinity.

*Effect of Ca^2+^/CaM on binding of R-synGAP to PDZ Domains of PSD-95–* We tested whether binding of Ca^2+^/CaM alters the binding of r-synGAP to PDZ domains by comparing binding to each affinity resin in the presence or absence of Ca^2+^/CaM (Fig. 8A). The presence of Ca^2+^/CaM during incubation with resin significantly reduces binding of r-synGAP to PDZ3 and to PDZ123, but not to PDZ1 and/or PDZ2. Thus, binding of Ca^2+^/CaM has a more specific, but weaker, effect on binding to the PDZ domains than does phosphorylation. The effects of phosphorylation and of the presence of Ca^2+^/CaM during incubation with resin are not additive (Fig. 8B); that is, the presence of Ca^2+^/CaM during the incubation with resin does not further decrease binding of phosphorylated r-synGAP to PDZ123.

**FIGURE 8.**
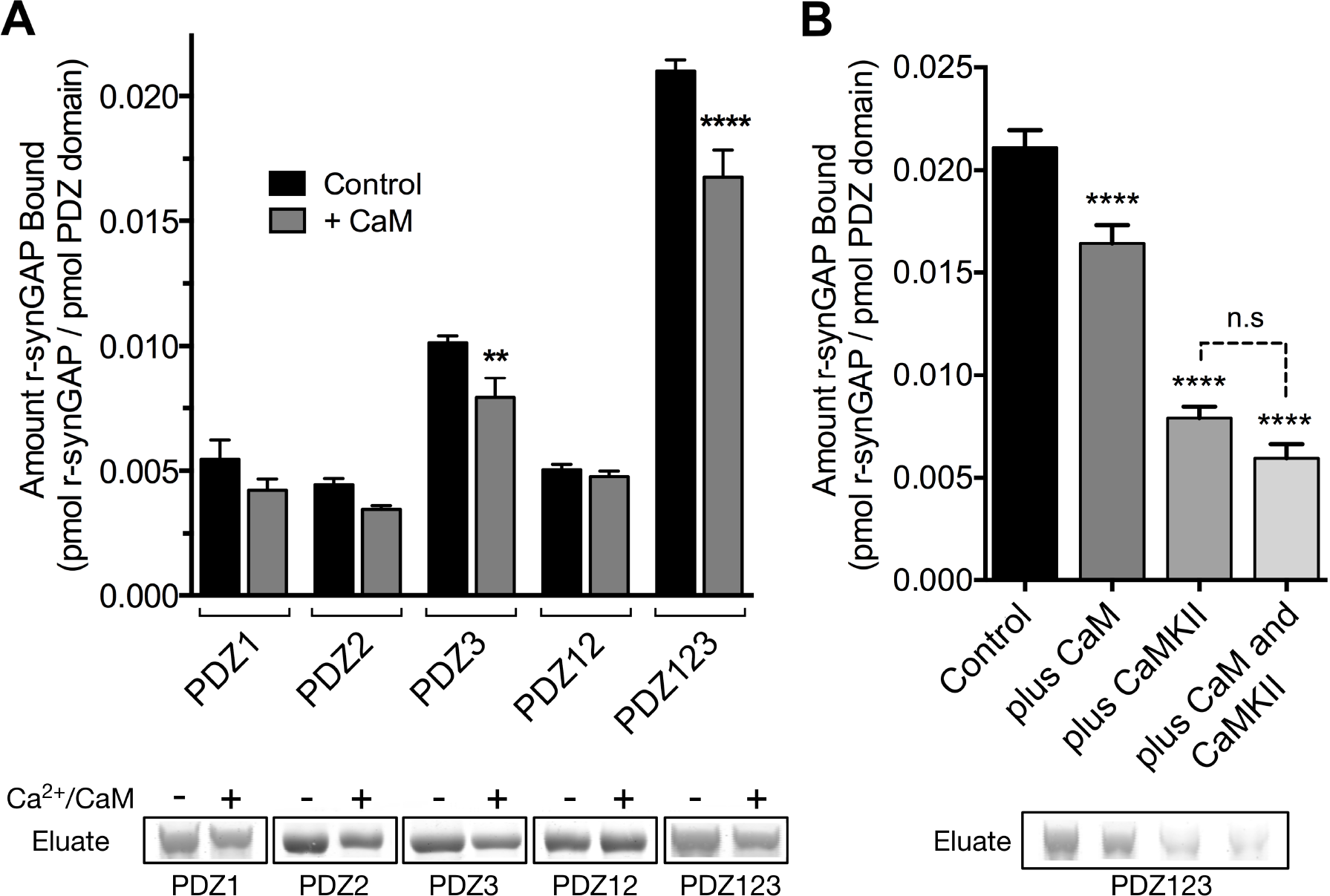
<u>Effect of Ca^2^/CaM binding on association of r-synGAP with PDZ domains of PSD-95.</u> *A,* Association of control and Ca^2+^/CaM bound r-synGAP with PDZ domains of PSD-95. R-synGAP (500 nM) without (Control) or with (+ CaM) 0.7 mM CaCl_2_/3.4 μM CaM was incubated with PDZ domain resins (PDZ1, PDZ2, PDZ3, PDZ12, and PDZ123) for 60 min at 25° C and bound synGAP was measured as described under Materials and Methods. For comparison of control to +Ca^2+^/CaM, PDZ1, p = 0.09; PDZ2, p = 0.2; PDZ3, p = 0.003 (d = 1.9); PDZ12, P = 0.7; PDZ123, p = 0.0001 (d = 2.6). *B,* Effects of bound Ca^2+^/CaM and phosphorylation by CaMKII on association of r-synGAP with PDZ123 domain are not additive. The association of synGAP with PDZ123 domain resin was measured as in A, under four different conditions: (Control), unphosphorylated r-synGAP alone; (plus CaM), Ca^2+^/CaM present in excess of synGAP during the incubation with resin; (plus CaMKII), r-synGAP is prephosphorylated with CaMKII as described under Materials and Methods, then Ca^2+^ is chelated with EGTA during the incubation with resin; and (plus CaM and CaMKII), r-synGAP is prephosphorylated by CaMKII and Ca^2+^/CaM is present in excess during incubation with resin. For comparison to Control: plus CaM, p<0.0001; plus CaMKII, p <0.0001; plus CaM and CaMKII, p <0.0001. For camparison of “plus CaMKII” and “plus CaM and CaMKII” samples, p = 0.06. Data are plotted as mean ± S.E. (n=4). The statistical significance of differences in PDZ domain binding relative to Control was determined by ordinary one way ANOVA (uncorrected Fisher's LSD). ^**^ *p*<0.01; ^****^ *p*<0.0001.

*SynGAP haploinsufficiency alters the composition of the PSD–* The physiological significance of the finding that phosphorylation by CaMKII decreases the affinity of synGAP for the PDZ domains of PSD-95 is best considered in the context of the high copy number of synGAP in the PSD. In molar terms, synGAP is approximately as abundant in the PSD as PSD-95 itself (Chen et al., 1998; Cheng et al., 2006; Dosemeci et al., 2007; Sheng and Hoogenraad, 2007). Because one synGAP molecule can bind to any one of the three PDZ domains in each molecule of PSD-95, synGAP could occupy as many as one third of the PDZ domains of PSD-95 in an average PSD. Our data suggest that phosphorylation of synGAP by CaMKII, triggered by activation of NMDARs, would promote dissociation of synGAP from the PDZ domains, reducing its ability to compete with other proteins for binding. We therefore propose that synGAP normally functions to limit the number of available PDZ domains in the PSD-95 complex, and that phosphorylation of synGAP by CaMKII partially alleviates the restriction, enabling reconfiguration of the PSD scaffold. This proposed function is distinct from synGAP’s role as a regulator of Ras and Rap and could explain its unusually high abundance in the PSD which, until now, has been mysterious (Chen et al., 1998; Sheng and Hoogenraad, 2007).

*SynGAP^−/+^* mice have been shown to have about half as much synGAP in homogenates from forebrain as *wild-type* litter mates. Because binding equilibria are driven not only by the intrinsic affinities of the binding partners, but also by their concentrations, one prediction of our proposed hypothesis is that synGAP haploinsufficiency, which reduces the amount of synGAP in the brain by 50% (Vazquez et al., 2004), will cause a significant increase in binding to PSD-95 of other prominent PSD-95 binding proteins, such as TARPs, LRRTM2s, or neuroligins. Thus, PSDs isolated from *synGAP^−/+^* mice would be predicted to have less synGAP and more TARPs, LRRTMs, and/or neuroligins bound to PSD-95 than do PSDs isolated from *wild-type* mice. We prepared PSD fractions from the forebrains of six *synGAP^−/+^* mice and from six *wild-type* litter mates and measured the ratios of synGAP, TARPs, LRRTM2, neuroligin-1, and neuroligin-2 to PSD-95 in the two fractions by quantitative immunoblot as described in Materials and Methods (Fig 9). As predicted, the level of synGAP is decreased in relation to PSD-95 by ~ 25% in PSDs (p = 0.0007) from the *synGAP^−/+^* mice (Fig. 9A). Furthermore, the ratios of TARPs 2,3,4,y8, and of LRRTM2 to PSD-95 are significantly increased (Fig. 9B,C; TARP/PSD-95, ~15%, p = 0.017; LRRTM2/PSD-95, ~16%, p = 0.0035). This result means that, as predicted, the increase in availability of PDZ1/2 domains on PSD-95 in the *synGAP^−/+^* mice enhances steady-state binding of TARPs and LRRTMs to those sites. Interestingly, the ratio of neuroligin-1 to PSD-95 is unchanged in the *synGAP^−/+^* mice (Fig. 9D), suggesting that increased availability of PDZ3 on PSD-95 is not a strong driver of association of neuroligin-1 with the PSD fraction. However, the ratio of neuroligin-2 to PSD-95 (Fig. 9E) is increased ~10% (p = 0.019). Neuroligin-2 normally associates mostly with inhibitory synapses and mediates their maturation (Varoqueaux et al., 2004—). However, Levinson et al. (2005) reported that over-expression of PSD-95 in neurons causes a redistribution of neuroligin-2, increasing the proportion associated with excitatory synapses. Taken together, these results verify the prediction that a decrease in availability of synGAP in the PSD scaffold, releases a restriction on binding of TARPs, LRRTMs, and neuroligin-2 to PSD-95 *in vivo* and increases their association with the PSD.

**FIGURE 9.**
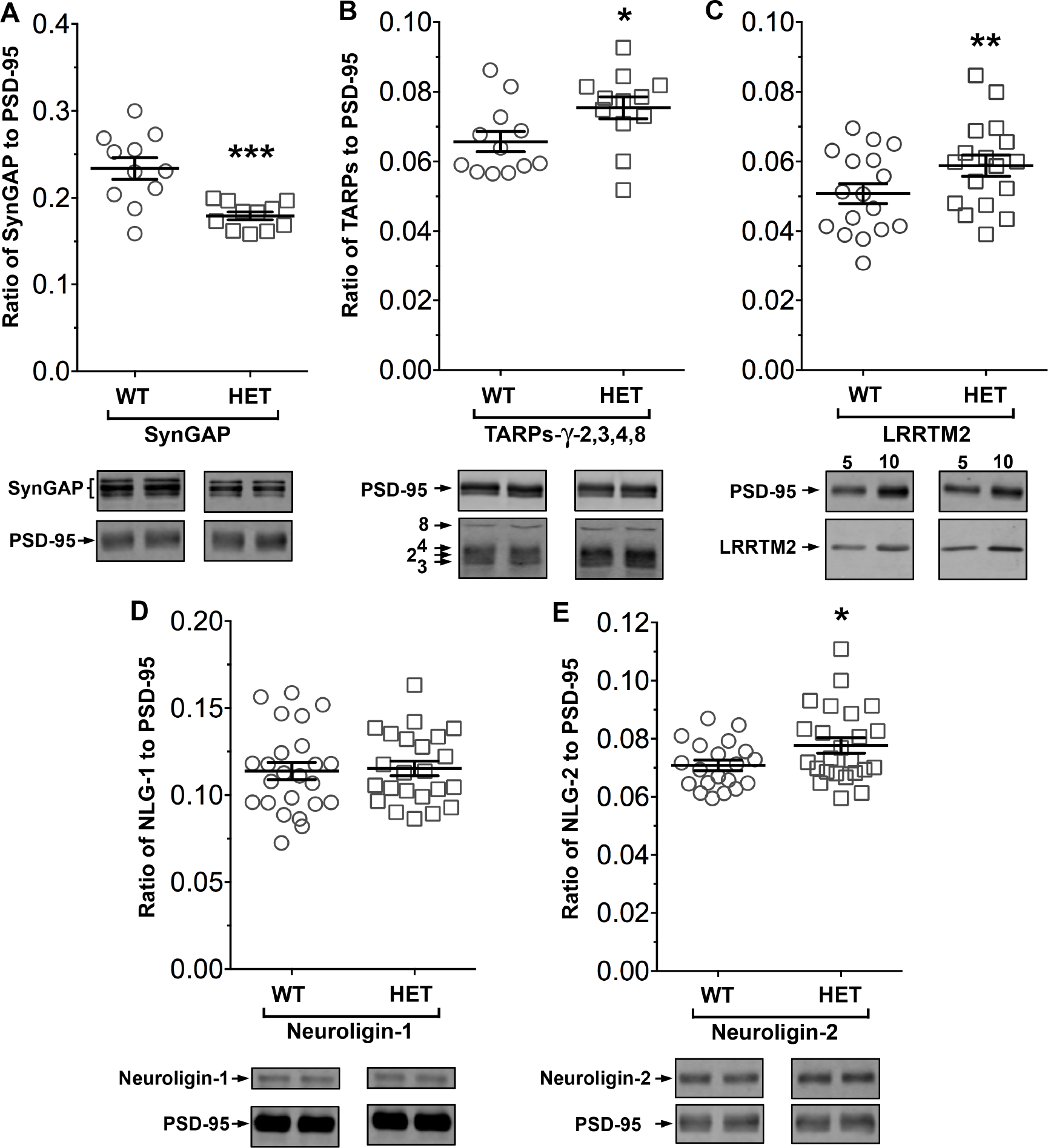
<u>Altered composition of the postsynaptic density in mice with heterozygous deletion of</u> <u>synGAP.</u> Ratios of amounts of the indicated proteins to PSD-95 were measured as described in Materials and Methods and are reported as mean ± S.E. For all blots except those for neuroligin-1, PSD-95 was detected with a secondary Ab labelled with AlexaFluor680 and the binding protein was detected with secondary Ab labelled with IRDye 800. On the neuroligin-1 blot, both PSD-95 and neuroligin-1 were detected with AlexaFluor680; the two bands were well-separated in each lane. Representative sets of visualized bands for *wild-type* (WT) and *synGAP^−/+^* (HET) from the same blot are shown below the graphs. A, SynGAP to PSD-95 ratio. Data were collected for 22 lanes from two blots containing 5 μg total PSD fraction per lane. One blot contained six lanes WT and six lanes HET samples, the other contained five lanes of each. The mean ratio of synGAP to PSD-95 was 0.234 ± 0.012 for WT (n = 11) and 0.179 ±0.005 (n = 11) for HET. Means were compared by unpaired, one-tailed t-test with Welch correction, p = 0.0007, d = 1.75. B, TARP y-2,3,4,8 to PSD-95 ratio. Data were collected for 24 lanes from two blots containing 10 μg total PSD fraction per lane. Each blot contained six lanes WT and six lanes HET samples. Densities of all four TARPs were pooled. The mean ratio of TARPs to PSD-95 was 0.066 ±0.003 (n = 12) for WT and 0.075 ±0.003 (n = 12) for HET. Means were compared by unpaired, one-tailed t-test with equal variance, p = 0.017, d = 0.93. C, LRRTM2 to PSD-95 ratio. Data were collected for 36 lanes from three blots containing six WT and six HET samples, alternating 5 and 10 μg (3 each). The mean ratio of LRRTM2 to PSD-95 was 0.051 ±0.003 for WT (n = 17) and 0.059 ±0.003 for HET (n = 17). Means were compared by paired, one-tailed t-test, p=0.0035, d = 0.66. D, Neuroligin-1 to PSD-95 ratio. Data were collected for 47 lanes from four blots two of which contained 5 μg and two 10 μg total PSD fraction per sample. Each blot contained six lanes WT and six lanes HET samples. The mean ratio of neuroligin-1 to PSD-95 was 0.114 ±0.005 (n = 24) for WT and 0.115 ± 0.004 (n = 23) for HET. Means were compared by unpaired one-tailed t-test, p = 0.413, d = 0.07. E, Neuroligin-2 to PSD-95 ratio. Data were collected for lanes from four blots containing 10 μg total PSD fraction per lane. Each blot contained six lanes WT and six lanes HET samples. The mean ratio of neuroligin-2 to PSD-95 was 0.071 ±0.002 (n = 20) for WT and 0.078 ± 0.003 (n = 24) for HET. Means were compared by unpaired, one-tailed t-test with Welch correction, p= 0.019, d = 0.64. *, p<0.05; ^**^, p<0.01; ^**^, p<0.001

## DISCUSSION

We have shown that phosphorylation of specific sites on synGAP by CaMKII substantially reduces the affinity of synGAP’s PDZ ligand for all three of the PDZ domains of PSD-95, and we hypothesize that this regulation helps to control the composition of the PSD at excitatory synapses. Recently, two other groups found that movement of synGAP out of the PSD can be triggered in living neurons by phosphorylation by CaMKII (Yang et al., 2013; Araki et al., 2015). In the Araki study, the authors suggested that phosphorylation of synGAP results in “dispersion” of synGAP away from the PSD and therefore would have the effect of upregulating Ras or Rap near the PSD. We propose here that a more important consequence of the decrease in binding of synGAP to the PDZ domains of PSD-95 is readjustment of the composition of the PSD resulting from increased availability of the PDZ domains of PSD-95. A cartoon version of this proposed mechanism is shown in Fig. 10. An average PSD (~360 nm in diameter) is estimated to contain ~300 molecules of PSD-95 with 900 PDZ domains (Chen et al., 2005; Sugiyama et al., 2005). Because synGAP is nearly as abundant in PSD-95 complexes as PSD-95 itself (Chen et al., 1998; Cheng et al., 2006; Dosemeci et al., 2007), synGAP could occupy as many as 300 PDZ domains in an average synapse. Other proteins that compete for binding to PDZ1 and 2 of PSD-95 include TARPs and LRRTMs. SynGAP may play an important role in limiting the size and strength of the synapse by limiting and helping to regulate the available “slots” that can bind AMPAR complexes (Hayashi et al., 2000; Shi et al., 2001; Opazo et al., 2012). Our hypothesis predicts that PSDs from mice with a deletion of one copy of the synGAP gene will contain fewer copies of synGAP and more copies of other proteins that bind to PSD-95. Indeed, we have shown here that PSD fractions from *synGAP^−/+^* mice have ~25% less synGAP per molecule of PSD-95 than those from *wild-type* mice; and, in contrast, they have significantly more TARP proteins (~15%), LRRTM2 (~16%), and neuroligin-2 (~9%) per molecule of PSD-95. The ratio of neuroligin-1 to PSD-95 is not altered.

**FIGURE 10.**
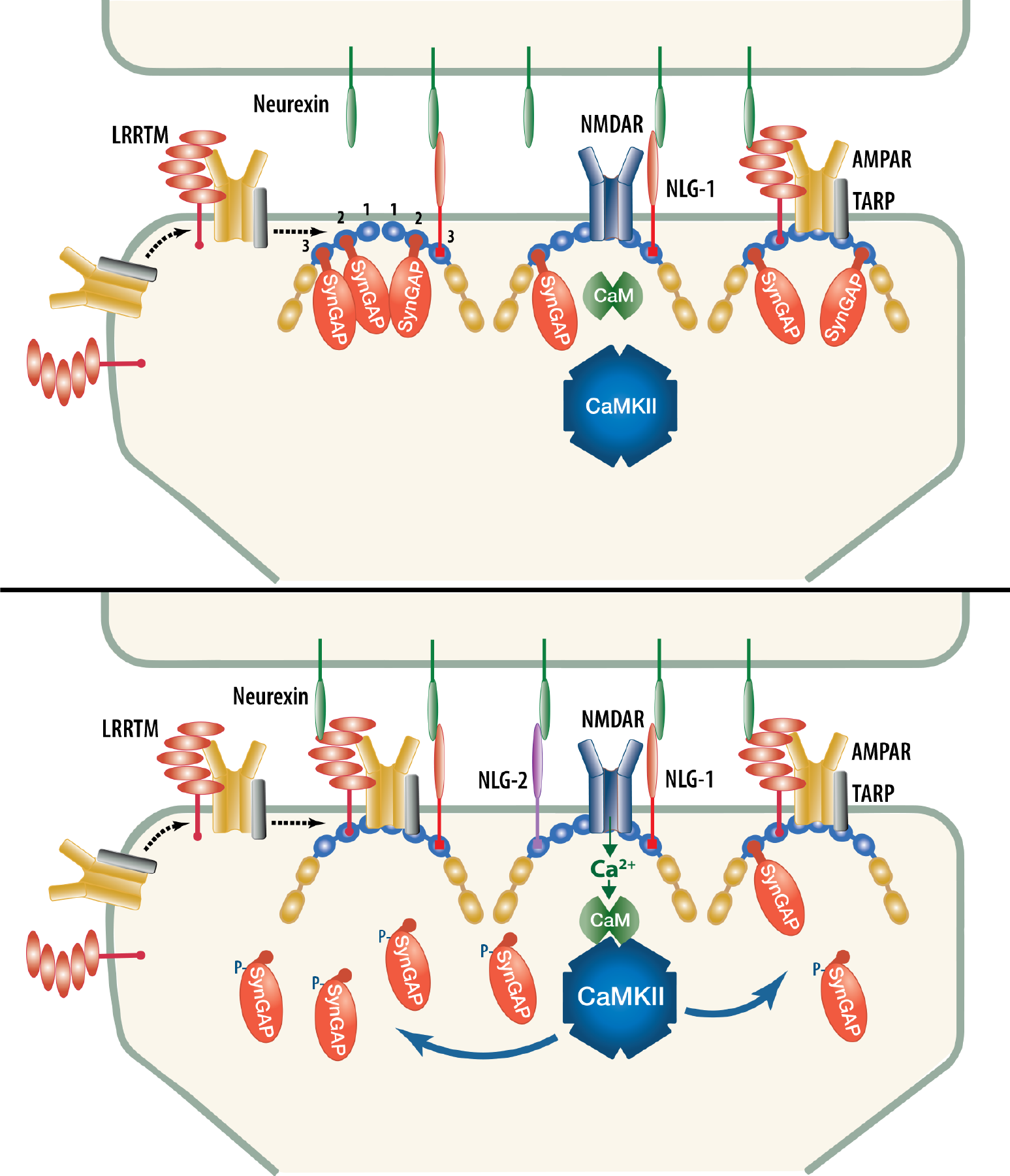
<u>Cartoon model of rearrangement of PSD after phosphorylation of synGAP by CaMKII.</u> *A,* Unphosphorylated synGAP binds to PDZ1, PDZ2 or PDZ3 of PSD-95, occupying as many as ~30% of its PDZ domains. The PDZ domains of PSD-95 are shown in blue and their numbers are indicated on the left pair of PSD-95 molecules. AMPARs that have been inserted into the extrasynaptic membrane by exocytosis associate with TARPs and with LRRTMs, both of which can bind to PDZ1 and PDZ2 of PSD-95. Neuroligin-1 (NLG1) binds to PDZ3 of PSD-95. LRRTMs and NLG1 also bind across the synaptic cleft to presynaptic neurexins. *B,* Calcium flux through NMDARs activates CaMKII leading to phosphorylation of synGAP on sites in the regulatory domain. The affinity of synGAP for the PDZ domains decreases, allowing TARPs, LRRTMs, and NLG-2 to displace synGAP by binding to the PDZ domains. The shift in affinity of synGAP creates “slots” that can be occupied by AMPAR complexes, or by neuroligin-2, leading to strengthening of the synapse.

Because binding between molecules is driven by both the inherent affinity between the binding components and by their concentrations, a reduction in binding of synGAP to PDZ domains on PSD-95 produced by phosphorylation by CaMKII, rather than by haploinsufficiency, will also lead to an increase in the amounts of TARP and LRRTM2, and/or neuroligin-2 in PSDs of *wild-type* animals. Acute phosphorylation of synGAP by CaMKII following activation of NMDARs during induction of LTP could initiate rearrangement of the composition of PSDs of individual synapses by causing an increase in equilibrium binding of TARPs, LRRTMs, and their associated AMPARs in the PSD as they diffuse from perisynaptic locations. Indeed, the dynamics of movements of synGAP and AMPARs visualized in living neurons following synaptic stimulation are consistent with this hypothesis. Activation of NMDARs and CaMKII causes dispersal of synGAP away from the PSD within a few minutes (Yang et al., 2013; Araki et al., 2015). The same stimuli produce an equally rapid increase in the rate of trapping of AMPARs at synaptic sites (Opazo et al., 2010; Opazo et al., 2012). Thus, the rates of these two processes observed in living neurons are compatible with the notion that reduced binding of synGAP to PSD-95 during induction of LTP opens up binding slots for AMPAR complexes.

Subsequent dephosphorylation of synGAP might be expected to allow synGAP to displace the newly added TARPs and LRRTMs. Thus, additional processes occurring later in the consolidation of LTP would be needed to be stabilize the newly added AMPARs in the synapse. These could include degradation of the phosphorylated synGAP and its replacement by newly synthesized alternatively-spliced isoforms that lack the C-terminal PDZ binding domain (McMahon et al., 2012).

The restriction by synGAP of binding of neuroligins, in particular neuroligin-2, to PDZ3 of PSD-95 may be more significant during development, during formation of new synapses, or perhaps in later phases of consolidation of LTP in adults. It is not clear whether a pool of perisynaptic neuroligins exists that could be readily recruited to new synaptic sites over a few minutes after phosphorylation of synGAP. In any case, the increased steady state amounts of TARPs, LRRTMs, and the small increase in neuroligin-2 that we observe in excitatory PSDs isolated from forebrain would increase the overall excitatory/inihibitory (E/I) balance of synapses onto neurons (Levinson and El-Husseini, 2005).

In addition to predicting the altered composition of the PSD in *synGAP^−/+^* mice, our hypothesis provides a mechanism to explain puzzling results from earlier studies. Neurons cultured from synGAP deficient mice have been reported by several groups to have a higher average number of AMPARs at their synapses than wild-type neurons (Kim et al., 2003; Vazquez et al., 2004; Rumbaugh et al., 2006). The two PSD-95 binding proteins that are proportionally increased in synGAP heterozygotes, TARPs and LRRTMs, bind subunits of AMPARs, and are believed to dock them in the synapse by binding to the PDZ domain “slots” on PSD-95 (Tomita et al., 2005; de Wit et al., 2009). Our data indicates that the increase in AMPARs in *synGAP+^/−^* mice is a direct result of increased binding of TARPs and LRRTM to PDZ domains that are made available by the reduced amount of synGAP.

A second example is the observation that absence of synGAP in hippocampal neurons cultured from synGAP deficient mice leads to accelerated maturation of spines, including early movement of PSD-95 into spine heads, and ultimately larger clusters of PSD-95 in individual synapses compared to *wild-type* neurons (Vazquez et al., 2004). We found that expression of *wild-type*-synGAP in the mutant neurons rescued all of these phenotypes; however, expression of synGAP with a deletion of the five residue PDZ ligand (*ΔSXV*) failed to rescue any of the effects of synGAP deficiency on accelerated maturation of spines. In fact, expression of synGAP-*ΔSXV* actually further increased the size of clusters of PSD-95 in spines compared to *wild-type* neurons. This failure to rescue the phenotypes was not a result of mislocalization of synGAP; synGAP-*ΔSXV* localized normally to developing spine heads. Instead, the data are consistent with the idea that synGAP normally competes with several proteins for binding to the PDZ domains of PSD-95, and thus limits the size of clusters of PSD-95 and its associated proteins, as well as their movement into spine heads (Vazquez et al., 2004).

We note that the reduction in affinity of synGAP for PDZ domains of PSD-95 after phosphorylation by CaMKII is apparently not sufficient for complete dispersal of synGAP away from the PSD, although it is likely necessary. We found that *synGAP-ΔSXV,* which cannot bind to PDZ domains, still localizes to synaptic spines (Vazquez et al., 2004). Furthermore, Barnett et al. (Barnett et al., 2006) showed that, in developing S1 rodent cortex, isoforms of synGAP that do not contain the PDZ ligand are, nevertheless, bound to the PSD. These data mean that reduction of affinity of synGAP for PDZ domains may be sufficient to decrease its ability to compete for binding slots on PSD-95; however, complete detachment of synGAP from the PSD *in vivo* appears to require additional events.

Others have proposed that, in general, PDZ domains act as flexible protein interaction points that can be modified to support changes in cytoplasmic organization (Kurakin et al., 2007). Complexes formed by PDZ domain interactions are examples of linked equilibria, the stable configurations of which are determined by the concentrations of each component and by their affinities for the relevant PDZ domains. Evidence has indicated that PSD-95 protein complexes exist in dynamic equilibrium permitting continual turnover and potential rearrangement of their composition (Sturgill et al., 2009; Schapitz et al., 2010). SynGAP is an abundant protein in the PSD with a relatively high affinity for all three of the PDZ domains of PSD-95; therefore, it will occupy a relatively large number of its PDZ domains at equilibrium. However, when synGAP’s affinities for the PDZ domains are reduced after phosphorylation, other components will begin to compete more effectively for binding and the composition of the PSD-95 complex will shift to a new equilibrium.

The functional significance of our finding that synGAP contains a high affinity binding site for Ca^2+^/CaM is less clear. We have shown that binding of Ca^2+^/CaM alters the conformation of the carboxyl terminal regulatory domain of synGAP allowing CDK5 to phosphorylate additional sites; the binding also reduces the affinity of synGAP for PDZ3 by ~25%. However, the consequences of these two effects for synaptic function are not known. However, once again, the high copy number of synGAP in the PSD may provide a clue. The biochemical events initiated by Ca^2+^ flux through NMDARs that lead to changes in synaptic strength (Sjostrom and Nelson, 2002) are initiated by formation of transient and limiting concentrations of Ca^2+^/CaM in the spine (Markram et al., 1998; Pepke et al., 2010). Approximately ten regulatory enzymes compete for binding of, and activation by, this Ca^2+^/CaM (Kennedy, 2013). Because of the abundance of synGAP in the PSD, the high affinity binding site for Ca^2+^/CaM on synGAP will compete effectively for the newly formed Ca^2+^/CaM and may act as a Ca^2+^/CaM buffer.

Haploinsufficiency of synGAP in humans is the cause of ~5% of cases of nonsyndromic cognitive disability and ~9% of such cases with co-morbid Autism Spectrum Disorder or Epilepsy (Berryer et al., 2013). The reduced amount of synGAP and resulting decrease in its ability to compete for PDZ domains are almost certainly significant factors in the pathology of these disorders, perhaps more significant than the reduction in synaptic Ras/Rap GAP activity. The increase in E/I balance in the forebrain predicted by our results could produce the symptoms of cognitive disability, ASD, and/or epilepsy observed in humans with synGAP haploinsufficiency. It would be worth considering whether pharmaceutical agents could be designed that would bind to PDZ domains of PSD-95 and compensate for the reduced level of synGAP.

## Materials and Methods

*Cloning, Expression and Purification of R-synGAP–* Soluble, recombinant synGAP (r-synGAP), comprising residues 103-1293 in synGAP A1-α1 (118-1308 in synGAP A2-α1), or sr-synGAP, comprising residues 103-725 in synGAP A1-α1 (118-740 in synGAP A2-α1), was purified from *E. coli* as previously described (Walkup IV et al., 2015). The isoform names and residue numbering are taken from ref. (Walkup IV et al., 2015). Henceforth, except where indicated, we use residue numbering corresponding to synGAP A1-α1.

Briefly, a pET-47b(+) plasmid (EMD Millipore, Billerica, MA, catalog no. 71461) containing synGAP cDNA (AF048976) fused to an N-terminal 6x Histidine Tag and a PreScission Protease cleavage site was transformed into the Rosetta2(DE3) *E. coli* strain (EMD Millipore, catalog no. 71397) for protein expression. Bacterial pellets were harvested by centrifugation and lysed by microfluidization in a ML-110 microfluidizer (Microfluidics Corp., Westwood, MA). Soluble synGAP was purified on Talon Metal

Affinity Resin (Clontech, Mountain View, CA, catalog no. 635503), and concentrated by ultrafiltration through a 150 kDa cutoff-filter (Thermo Scientific, Waltham, MA, catalog no. 89923) for r-synGAP or 9 kDa cutoff-filter (Thermo Scientific, catalog no. 89885A) for sr-synGAP. Concentrated samples of r-synGAP were exchanged into storage buffer (20 mM Tris, pH 7.0; 500 mM NaCl, 10 mM TCEP, 5 mM MgCl_2_, 1 mM PMSF, 0.2% Tergitol Type NP-40, and Complete EDTA-free protease inhibitor) by ultrafiltration, flash-frozen in liquid nitrogen, and stored at-80° C. Sr-synGAP was further purified on a size exclusion column prior to storage (Walkup IV et al., 2015).

*Cloning, Expression and Purification of PDZ Domains from PSD-95–* Soluble recombinant PDZ domains, comprising residues 61-151 (PDZ1),155-249 (PDZ2), 302-402 (PDZ3), 61-249 (PDZ12), and 61-403 (PDZ123) from murine PSD-95 (Q62108) were purified from *E. coli* as previously described (Walkup IV and Kennedy, 2014, 2015) with the modifications below.

Briefly, pJExpress414 plasmids (DNA2.0, Menlo Park, CA, catalog no. pJ414) containing codon optimized PDZ domains were transformed into the BL21(DE3) *E. coli* strain (EMD Millipore, catalog no. 70235-3) for protein expression. Single colonies of BL21(DE3) cells harboring pJExpress414 plasmids were grown overnight at 37 °C in lysogeny broth (LB) (Teknova, Hollister, CA, catalog no. L9110) supplemented with 100 μg/ml carbenicillin. Overnight cultures were diluted 1:500 into LB medium and grown at 37 °C until cultures reached an O.D._600_ of 1.0. IPTG was added to a final concentration of 0.2 mM, and cultures were grown for an additional 4.5 hours at 37 °C. Bacterial pellets were harvested by centrifugation and lysed using non-ionic detergent (BugBuster) and ReadyLyse. Soluble PDZ1, PDZ2 and PDZ12 domains were purified on GluN2B peptide (GAGSSIESDV) PDZ Ligand Affinity Resin (Walkup IV and Kennedy, 2014) by eluting with 400 μg/ml SIETEV peptide. PDZ3 and PDZ123 were purified on CRIPT peptide (GAGNYKQTSV) PDZ Ligand Affinity Resin (Walkup IV and Kennedy, 2014) by eluting with 400 μg/ml YKQTSV peptide. PDZ domains were concentrated by ultrafiltration through a 3 kDa Amicon Ultracentrifugal Filter Unit (EMD Millipore, catalog no. UFC 900396). The PDZ peptide ligands were removed from PDZ domains by dialysis into storage buffer (50 mM HEPES, pH 7.5; 100 mM NaCl, 5 mM TCEP, 1 mM PMSF, and Complete EDTA-free protease inhibitor). Purified PDZ domains (>99% pure; 45-610 μM; Fig. 1 figure supplement 1) were flash-frozen in liquid nitrogen, and stored at-80° C.

**FIGURE 1-figure supplement 1.**
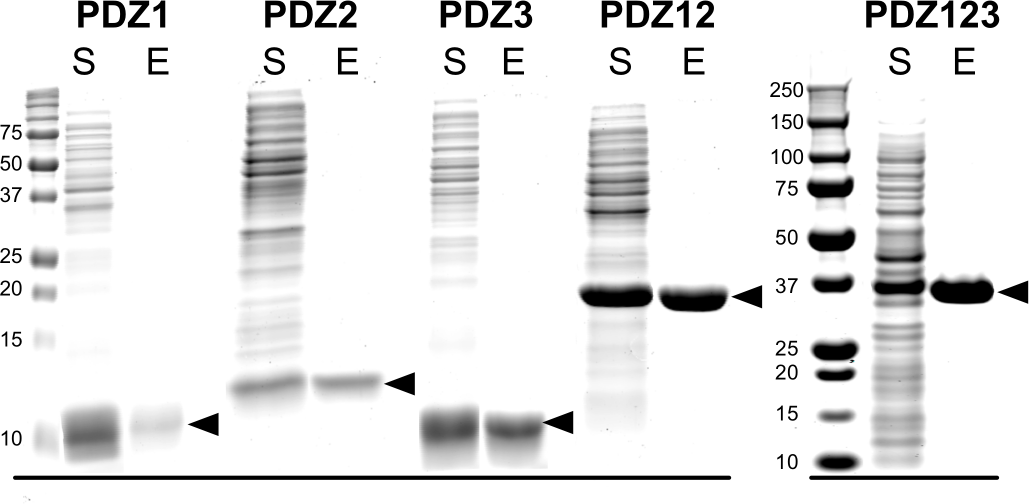
<u>Purification of recombinant PDZ domains of PSD-95.</u> PDZ domains of PSD-95 were expressed in *E. coli* individually or combined in a single peptide, as indicated, and purified as described under “Materials and Methods.” Proteins in the starting soluble fraction (S) and eluted from PDZ ligand affinity columns (E) were fractionated on 12% (PDZ1, PDZ2, PDZ3, PDZ12) or 4-12% gradient (PDZ123) SDS-polyacrylamide gels and stained as described under “Materials and Methods.”

*SDS-PAGE, Immunoblotting and Assessment of Protein Purity and Yield–* We used SDS-PAGE to determine purity of proteins and to quantify binding to PDZ domain resin. Protein samples were diluted 1:3 into 3x Laemmli buffer (100 mM Tris HCl, pH 6.8; 2.1% SDS, 26% glycerol, 7.5% P-mercaptoethanol, and 0.01% bromophenol blue) and heated to 95° C for 3 minutes before fractionation on 8% SDS-PAGE gels at 165 V in 25 mM Tris base, 192 mM glycine, 0.1% SDS. Proteins were stained with Gel Code Blue (Thermo Scientific, catalog no. 24592), imaged on a LI-COR Odyssey Classic Infrared Imaging System (LI-COR Biosciences, Lincoln, NE) at 700 nm, and quantified with LI-COR Image Studio Software (v4.0.21) against standard curves of BSA (catalog no. A7517-1VL) and lysozyme (catalog no. L4631-1VL) purchased from Sigma-Aldrich, St. Louis, MO. The protein standards were loaded onto each gel in lanes adjacent to the protein samples. Molecular weights of stained proteins were verified by comparison to Precision Plus Protein All Blue Standards (BioRad, Hercules, CA, catalog no. 161-0373).

For immunoblotting, proteins fractionated by SDS-PAGE were electrically transferred to low fluorescence PVDF membranes (Thermo Scientific, catalog no. 22860) in 25 mM Tris, 200 mM glycine, and 20% methanol. Membranes were washed with 50 mM Tris-HCl, pH 7.6; 150 mM NaCl (TBS) followed by blocking with Odyssey Blocking Buffer (LI-COR Biosciences, catalog no. 927-50000). Membranes were washed in TBS supplemented with 0.1% Tween 20 (TBS-T) before incubation in Odyssey Blocking Buffer containing 1:1000 diluted rabbit anti-synGAP (Pierce, Rockford, IL, catalog no. PA1-046) or 1:1500 BSA-free anti-TetraHis (Qiagen, Germany, catalog no. 34670). Bound antibodies were detected with 1:10,000 goat anti-mouse Alexa-Fluor 680 (Life Technologies, Carlsbad, CA, catalog no. A-21057) or 1:10,000 goat anti-rabbit Alexa-Fluor 680 (Life Technologies, catalog no. A-21109) visualized with a LI-COR Odyssey Classic Infrared Imaging System and quantified with LI-COR Image Studio Software.

*Synthesis of PDZ Domain Affinity Resins–* PDZ domain affinity resins (PDZ1, residues 61-151; PDZ2, residues 155-249; PDZ3, residues 302-402; PDZ12, residues 61-249; PDZ123, residues 61-402 from murine PSD-95) were prepared by the HaloTag-HaloLink method as previously described (Walkup IV and Kennedy, 2014, 2015). Briefly, bacterial cell pellets containing PDZ domain-HaloTag fusion proteins were resuspended in 10 ml/g of Purification Buffer, and lysed by three passes through a ML-110 microfluidizer. The lysate was clarified by centrifugation, added to HaloLink resin (Promega, Madison, WI, catalog no. G1915), and mixed with continuous agitation for 1.5 hours at 4° C on an end-over-end mixer. Unbound protein was separated from the resin by centrifugation and the PDZ-HaloTag-HaloLink resin was resuspended, transferred to a column, and allowed to settle. The resin was extensively washed and then stored at 4° C in a buffer supplemented with 0.05% NaN_3_. The resin was used or discarded within 1 week of preparation. The densities of PDZ domains on the resin varied from 50 to 100 pmol of PDZ123 domain and from 200 to 500 pmol of PDZ1, PDZ2, PDZ3, or PDZ12 domains per μl resin.

*Assay for Binding of R-synGAP to PDZ domain Resin–* Phosphorylated or non-phosphorylated synGAP (500 nM, 200 μl) was mixed with 20 μl of affinity resin containing PDZ1, PDZ2, PDZ3, PDZ12, or PDZ123 domains, pre-equilibrated with Binding/Wash Buffer (25 mM Tris, pH 7.0; 150 mM NaCl, 1 mM MgCl_2_, 0.5 mM TCEP, 0.2% Tergitol, 0.5 mM EDTA) in a cellulose acetate spin cup (Pierce, catalog no. 69702) for 60 min on an end-over-end mixer. In some experiments, 2.5 μM CaM and 0.5 mM CaCl_2_ were included to test the effect of binding of Ca^2+^/CaM to synGAP on synGAP’s affinity for PDZ domains. After the incubation, the resin in the spin cup was centrifuged for 2 min at 1,500 × g to remove unbound protein, and the resin was washed 4 times with 200 μl of Binding/Wash Buffer. To elute bound protein, 100 μl of 1× Laemmli Buffer was added and the resin was incubated for 5 min at room temperature. The eluted protein was collected by centrifugation at 6,000 × g for 2 min, fractionated by SDS-PAGE, stained with Gel Code Blue, and quantified on a LI-COR Classic as described above.
Integrated intensities reflecting the amount of bound synGAP were determined with LI-COR software and plotted with Prism (v6.0d, GraphPad Software, La Jolla CA).

*Phosphorylation of R-synGAP by CaMKII for PDZ Binding Assays–* Phosphorylation of r-synGAP by CaMKII was carried out immediately prior to PDZ binding assays, as previously described (Walkup IV et al., 2015). Reaction mixtures contained 50 mM Tris-HCl, pH 8.0; 10 mM MgCl_2_, 0 or 0.7 mM CaCl_2_, 0.4 mM EGTA, 30 μM ATP, 0 or 3.38 μM CaM, 10 mM DTT, 725 nM r-synGAP, and 0 or 10 nM CaMKII. Samples were quenched by addition of 1/3 volume of 50 mM Tris, pH 8.0; 0.4 M NaCl, 10 mM MgCl_2_, 0.8% tergitol (Type NP-40), 6 μM autocamtide-2 related inhibitory peptide (Genscript, China), 40 mM EGTA at the indicated times. When we planned to add Ca^2+^/CaM during the subsequent incubation with resin, the EGTA was omitted. Samples were stored on ice until their use in PDZ domain binding assays.

*Stoichiometry and Rate of Phosphorylation of R-synGAP and Histone H1 by CaMKII and/or CDK5–* Phosphorylation of 725 nM synGAP by 10 nM CaMKII was carried out as previously described (Walkup IV et al., 2015) in reaction mixtures containing 50 mM Tris-HCl, pH 8.0, 10 mM MgCl_2_, 0 or 0.7 mM CaCl_2_, 0.4 mM EGTA, 500 μM [γ-^32^P]-ATP (375 cpm/pmol) (6000 Ci/mmol, Perkin Elmer, Waltham, MA, catalog no. BLU002Z/NEG002Z), 0 or 3.4 μM CaM, 10 mM DTT. Phosphorylation of 286 nM r-synGAP and 4.3 μM Histone H1 (New England Biolabs, Ipswich, MA, catalog no. M2501S), by CDK5/p35 (EMD Millipore, catalog no. 14-477M) or CDK5/p25 (EMD Millipore, catalog no. 14-516) was carried out in the same reaction mixture containing 110 nM CDK5/p35 or CDK5/p25 but no CaMKII. After fractionation by SDS-PAGE, phosphorylated proteins were quantified with a Typhoon LA 9000 phosphorimager (GE Healthcare Life Sciences, Pittsburg, PA) as previously described (Walkup IV et al., 2015). Relative densities were converted to pmol phosphate by comparison to densities of standard amounts of [γ-^32^P]-ATP. The stoichiometry of phosphorylation was calculated by dividing mol of incorporated phosphate by mol of synGAP loaded per lane.

*Measurement of Affinity of R-synGAP for PDZ Domains by Surface Plasmon Resonance (SPR)–* We used a “competition in solution” method (also called “affinity in solution”) (Nieba et al., 1996; Lazar et al., 2006; Abdiche et al., 2008) to measure the affinity of r-synGAP for PDZ domains. In this method, PDZ domains are immobilized on the chip surface and used to capture and measure the concentration of free r-synGAP in pre-equilibrated mixtures of a constant amount of r-synGAP with varying amounts of soluble recombinant PDZ domains. Experiments were performed on a Biacore T200 (GE Healthcare Life Sciences). Purified PDZ domains (PDZ1, PDZ2, PDZ3, PDZ12, PDZ123 from PSD-95) were coupled to Series S CM5 Sensor Chips (GE Healthcare Life Sciences, catalog no. BR-1005-30) by the amine coupling protocol specified in the Biacore T200 Control Software with reagents purchased from GE Healthcare Life Sciences. Sensor surfaces were activated by applying a 1:1 mixture of 50 mM N-hydroxysuccinimide (NHS): 200 mM 1-ethyl-3-(3-dimethylaminopropyl) carbodiimide hydrochloride (EDC) provided in the Biacore Amine Coupling Kit (GE Healthcare Life Sciences, catalog no. BR-1000-50) dissolved in HBS-N running buffer (degassed 0.01 M HEPES pH 7.4, 0.15 M NaCl) (GE Healthcare Life Sciences, catalog no. BR-1006-70). PDZ domains were diluted to 0.1–5 μM in Biacore Sodium Acetate Buffer [10 mM sodium acetate, pH 4.0 (GE Healthcare Life Sciences, catalog no. BR-1003-49) for PDZ1 and PDZ3; pH 4.5 for PDZ2; pH 5 for PDZ12 and PDZ123]. PDZ domains were injected into flow cells 2 and 4 until 200 to 400 RU (resonance units) of PDZ domain were immobilized. Flow cells 1 and 3 were left blank to be used as reference surfaces. Ethanolamine (1 M, pH 8.5) was injected for 7 min at 10 μl/min to block remaining active sites on all four flow cells.

A calibration curve was prepared by applying samples of 0 to 50 nM r-synGAP prepared by two-fold serial dilution of 50 nM r-synGAP into 1× HBS-EP+ buffer (degassed 0.01 M HEPES, pH 7.4; 0.15 M NaCl, 3 mM EDTA, 0.05% v/v Surfactant P20; GE Healthcare Life Sciences, catalog no. BR-1006-69) to the chip and recording the maximum RU for each concentration. Samples for calibration were incubated for 2 hours at room temperature before randomized injection onto the chip surface at 25° C at 10 μl/min for 200 sec over all four flow cells. Between each sample injection, the chip was regenerated by injecting 1 M MgCl_2_ at 100 μl/min for 60 sec, waiting 180 sec for the baseline to stabilize, then injecting a second pulse of MgCl_2_ solution, waiting 300 sec for the baseline to stabilize, and finally executing a “carry over control injection” in which HBS-EP+ buffer is flowed over the chip surface at 40 μl/min for 30 sec.

Mixtures of r-synGAP and PDZ domains were prepared by 1:1 dilution of 50 nM r-synGAP with twofold serial dilutions of PDZ domains (0-20 μM PDZ1, PDZ2, PDZ3, PDZ12 or 0-160 μM PDZ123) in HBS-EP+ buffer to produce mixtures containing 25 nM r-synGAP and 0-10 μM PDZ1, PDZ2, PDZ3, PDZ12 or 0-80 μM PDZ123. For each mixture of r-synGAP and PDZ domain, the different concentrations were injected randomly and a series of sensorgrams were recorded as described for the calibration curve.

Sensorgrams were processed with Biacore T200 Evaluation Software, (ver. 3.0, GE Healthcare Life Sciences). The y-axes were zeroed at the baseline for each cycle and x-axes were aligned at the injection start. Bulk refractive index changes and systematic deviations in sensorgrams were removed by subtracting the responses in reference flow cells corresponding to the sample flow cells (e.g. 2-1, 4-3). The averaged sensorgrams for 0 nM r-synGAP were then subtracted from sensorgrams for all other concentrations. The concentrations of free r-synGAP in each mixture with PDZ domains was determined from the calibration curve, exported into Prism, and plotted against the log of PDZ domain concentration. The curve was fit to the equation:

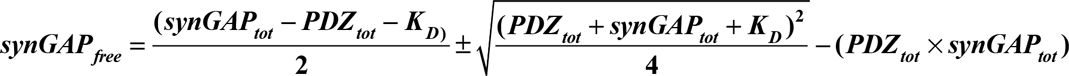

Equilibrium dissociation constants (K_D_) for binding to PDZ1, PDZ2, and PDZ3, and apparent equilibium dissociation constants for binding to PDZ12 and PDZ123 (K_Dapp_) were determined from the best fit curves as described in the Biacore T200 Software Handbook (p. 210).

*Binding of R-synGAP to CaM-Sepharose–* Rosetta2(DE3) cells containing sr-synGAP or r-synGAP were lysed in Lysis buffer as described in Walkup et al. (2015) except that the buffer contained 200 mM NaCl, 0 or 5 mM CaCl_2_, and 0 or 10 mM EGTA. The resuspended cells were lysed by sonication with a Digital Sonifier 450 Cell Disruptor (Branson, Danbury, CT) for two passes at 90 seconds/pass (15% power, 1.0 second on, 1.5 seconds off), and insoluble material was removed by centrifugation at 16,000 × g for 40 min at 4° C. Clarified cell lysate (1.7 ml) containing sr-synGAP or r-synGAP (~6 mg/ml total protein) was incubated with end-over-end mixing for 60 min with 0.3 ml CaM-Sepharose 4B (GE Healthcare Life Sciences, catalog no. 17-0529-01) or control Sepharose 4B (GE Healthcare Life Sciences, catalog no. 17-0120-01). The resin was pipetted into a BioSpin column (Bio-Rad, catalog no. 732-6008) and washed with 12 ml (40 column volumes) of Lysis/Wash Buffer. Bound protein was eluted with 1.2 ml (4 column volumes) of Lysis/Wash Buffer containing 100 mM EGTA. Eluted proteins (30 μl aliquot) were resolved by SDS-PAGE and transferred to a PVDF membrane which was probed with anti-synGAP or BSA-free anti-TetraHis as described above.

*Measurement of Affinity of CaM for R-synGAP by SPR–* Direct binding of r-synGAP to Ca^2^ /CaM immobilized on a chip was assayed on a Biacore T200 with a Series S Sensor Chip CM5. CaM (Enzo, Farmingdale, NY, catalog no. BML-SE325-0001) was coupled to the chip by the amine coupling protocol specified in the Biacore T200 Control Software, as described above. Purified, lyophilized CaM (250 μg) was resuspended in water and exchanged into Biacore Sodium Acetate, pH 4.0 Buffer (10 mM sodium acetate, pH 4.0) with an Amicon Ultra-0.5 ml centrifugal filter with a 3 kDa molecular weight cutoff (EMD Millipore, catalog no. UFC500396). CaM was further diluted to 0.5 nM in 10 mM Sodium Acetate, pH 4.0, and injected into flow cells 2 and 4 until 50 RU of CaM were immobilized (~7 min at a flow rate of 10 μl/min). Flow cells 1 and 3 were left blank to be used as reference surfaces. Ethanolamine (1 M, pH 8.5) was injected for 7 min at 10 μl/min to block remaining active sites on all four flow cells. R-synGAP (0 nM to 75 nM) in 1× HBS-EP+ running buffer supplemented with 10 mM CaCl_2_, was injected in triplicate at 25° C at 100 μl/min for 75 sec over all four flow cells. Different concentrations of r-synGAP were applied in randomized order. After injection ended, dissociation was monitored in each flow cell for 500 sec. Regeneration of the sensor chip was performed by injecting 50 mM NaOH (GE Healthcare Life Sciences, catalog no. BR-1003-58) at 100 μl/min for 30 sec, waiting 180 sec for the baseline to stabilize, then injecting a second pulse of NaOH, waiting 240 second for the baseline to stabilize, and finally executing a “carry over control injection.” Sensorgrams were processed using the Biacore T200 Evaluation Software, version 3.0, as described above. Resonance units of bound synGAP at equilibrium were exported into Prism and plotted against the concentrations of r-synGAP. The data were fit globally to a hyperbolic curve by nonlinear regression to determine equilibrium dissociation constants (Kd).

*Determination of Affinity of Ca^2+^/CaMfor Sr-and R-synGAP by Equilibrium Binding to CaM-Sepharose–* Purified sr-synGAP and r-synGAP were diluted to 1 to 500 nM in Binding/Wash Buffer (50 mM Tris, pH 7.5; 200 mM NaCl, 5 mM TCEP, 2 mM CaCl_2_). Aliquots of diluted synGAP (300 μl) were incubated with end-over-end mixing for 60 minutes with 50 μl of CaM-Sepharose 4B in a screw cap spin column (Thermo Scientific, catalog no. 69705). Concentrations of sr-and r-synGAP were 20-3,000 fold below the ligand binding capacity of the CaM-Sepharose resin. Unbound protein was removed by centrifugation at 4,000 × g for 30 seconds. Beads were washed in Binding/Wash Buffer (250 μl, 5 volumes), and bound protein was eluted with 50 μl of 1× Laemmli buffer with 10% β-mercaptoethanol. Eluted proteins were fractionated by SDS-PAGE and transferred to a PVDF membrane as described above. Blots were probed with 1:1000 anti-synGAP or 1:1500 BSA-free anti-TetraHis anti-bodies and quantified on a LI-COR Classic as described above. Integrated intensities reflecting the amount of bound synGAP were determined with LI-COR software and plotted against the corresponding concentrations of synGAP with Prism software. The data were fit to a hyperbolic curve by nonlinear regression to determine the dissociation constant (K_D_).

*Preparation of PSD fractions–* PSD fractions were prepared as previously described (Cho et al., 1992) from six *wild-type* and six *SynGAP^−/+^* mouse litter mates matched by age (7-12 weeks), and sex *(wild-type,* 1 female, 5 males; *synGAP^−/+^,* 2 female, 4 male). The mice were killed by cervical dislocation and forebrains were dissected and rinsed in Buffer A (0.32 M sucrose, 1 mM NaHCO3, 1 mM MgCl_2_, 0.5 mM CaCl_2_, 0.1 mM PMSF, 1 mg/l leupeptin). Forebrains from each set of six mice were pooled and homogenized with 12 up and down strokes at 900 rpm in 14 ml Buffer A. Homogenates were diluted to 35 ml in Buffer A and centrifuged at 1400 × g for 10 min. The pellet was resuspended in 35 ml Buffer A, homogenized (3 strokes), and centrifuged at 710 g for 10 min. Supernatants from the two centrifugations were combined and centrifuged at 13,800 g for 10 min. The pellet was resuspended in 8 ml of Buffer B (0.32 M sucrose, 1 mM NaH_2_CO_3_), homogenized with 6 strokes and layered onto a sucrose gradient (10 ml each of 0.85 M, 1.0 M, and 1.2 M sucrose in 1 mM NaH_2_CO_3_ buffer). The gradient was centrifuged for 2 hours at 82,500 g in a swinging bucket rotor. The synaptosome-enriched layer at the interface of 1.0 and 1.2 M sucrose was collected, diluted to 15 ml with Solution B and added to an equal volume of Buffer B containing 1% Triton. The mixture was stirred for 15 min at 4°C and centrifuged for 45 min at 36,800 g. The pellet containing the PSD-enriched, Triton-insoluble fraction was resuspended in 800-1000 μl of 40 mM Tris pH 8 with a 21 gauge needle and 1 ml syringe, and further solubilized by hand in a teflon glass homogenizer. Samples were aliquoted and stored at-80° C.

*Quantification of Proteins in PSD Fractions–* PSD samples contained six pooled biological replicates of each of the two genotypes. Equal amounts of protein from each PSD sample (5-15 μg) were dissolved in SDS-PAGE sample buffer (33 mM Tris HCl, pH 6.8; 0.7% SDS, 10% glycerol, 2.5% β-mercaptoethanol, and 0.003% bromophenol blue), heated at 90 °C for 5 min, fractionated on polyacrylamide gels (8% or 10%), and electrically transferred to PVDF membranes in pre-cooled 25 mM Tris, 150 mM glycine, 2% methanol at 250V for 2.5 hours at 4°C. Membranes were blocked with Odyssey blocking buffer (LI-COR Biosciences) and then incubated in a primary antibody solution of 5% BSA in TBS-T overnight at 4°C, as described above. Primary antibodies included mouse-anti-PSD-95 (ThermoFisher, catalog no. MA1-046 [clone 7E3-1B8], dilution 1:10,000), rabbit-anti-SynGAP (Pierce, catalog no. PA1-046, dilution 1:3500), rabbit-anti-TARP (y-2,3,4, and 8; EMD Millipore, catalog no. Ab9876; dilution 1:300), rabbit-anti-LRRTM2 (Pierce, catalog no. PA521097; dilution 1:3000), mouse-anti-neuroligin-1 (Sigma, catalog no. sab5201464; dilution 1:250), and rabbit-anti-neuroligin-2 (Synaptic Systems, Germany, catalog no. 129202; dilution 1:1000). The membranes were then washed 3-times in TBS-T. The membrane was incubated with secondary antibodies (Alexa Fluor 680-goat-anti-mouse IgG (Life Technologies, catalog no. A21057; 1:10,000) or IRDye800-goat-anti-rabbit IgG (Rockland, Limerick, PA, catalog no. 611-132-122; 1:10,000) for 45 minutes at room temperature in 5% nonfat milk in TBS-T, then washed 3 times in TBS-T, then twice in TBS prior to scanning. For most experiments, each blot contained 6 duplicate samples of PSD fractions from *wild-type* and the same number from *synGAP^−/+^* mice. Each blot was incubated with a mixture of two primary antibodies; mouse-anti-PSD-95 and the antibody against neuroligin-1, neuroligin-2, TARPs, LRRTM2, or synGAP. Then the blots were incubated with a mixture of the appropriate secondary antibodies. For measurement of neuroligin-1, both PSD-95 and neuroligin-1 were detected by the same goat anti-mouse secondary; the bands were physically separated on the gel and were quantified independently. Bound antibodies were visualized in the appropriate fluorescent channels with an Odyssey Classic Infared Imaging System (LI-COR Biosciences, Lincoln, Nebraska).

Before running samples for quantification, we determined empirically the amount of PSD sample that would result in signals for each protein that were strong enough for measurement and not saturated. To quantify the densities of the bands, each visual image was first set to high brightness in order to capture the boundaries of the signals for each band. The images were then used as a template in LI-COR software to draw rectangular regions of interest around protein bands, and around identically sized background regions in the same lane. Background densities were subtracted from each protein signal. Two blots were excluded because background density formed a gradient across the gel resulting in more variation in measurement for one of the genotypes. The Digital data read by the LI-COR software is unchanged by the visualization settings and is linear over several orders of magnitude. For each lane, the ratio of the integrated density of each of the five proteins to the integrated density of PSD-95 was calculated. For three gels, one outlier measurement (defined as greater than 2 times the standard deviation of the mean) was excluded from the calculation. A few bands were also excluded from measurement when a bubble during the transfer distorted and blurred the signal. The mean and standard error of the mean (S.E) of the ratios were determined for *wild-type* and *synGAP^−/+^* PSD fractions. The means were compared by t-tests performed with Prism software as indicated in the legend of Fig. 9. For four of the proteins, the number of measurements was sufficient to determine a significant difference between *wild-type* and *synGAP^−/+^.* In the case of neuroligin-1, the means were identical after 24 individual measurements of each sample.

## Acknowledgements

This work was supported in part by grants from the Gordon and Betty Moore Foundation (Center for Integrative Study of Cell Regulation), the Hicks Foundation for Alzheimer's Research, the Allen and Lenabelle Davis Foundation, and from National Institutes of Health Grant MH095095 to MBK. WGW IV was supported by the National Science Foundation Graduate Research Fellowship under Grant No. 2006019582, and the National Institutes of Health under Grant No. NIH/NRSA 5 T32 GM07616. The Protein Expression Center is supported by the Beckman Institute.

**Conflict of interest**: The authors declare that they have no conflicts of interest concerning the contents of this article.

